# Pre-rRNA spatial distribution and functional organization of the nucleolus

**DOI:** 10.1101/2024.11.02.621631

**Authors:** Yu-Hang Pan, Lin Shan, Zheng-Hu Yang, Yu-Yao Zhang, Yuan Zhang, Shi-Meng Cao, Jun Zhang, Li Yang, Ling-Ling Chen

## Abstract

The self-organized nucleolus is where the pre-ribosomal RNA (pre-rRNA) processing and pre-ribosomal ribonucleoprotein (RNP) assembly take place. Here we present the spatiotemporal distribution of pre-rRNA intermediates in the human nucleolus, revealing a segregated spatial distribution of small subunit (SSU) and large subunit (LSU) precursors. Notably, the 5’ external transcribed spacer (5’ ETS)-containing SSU pre-rRNAs are retained across FC-DFC-PDFC regions, while the internal transcribed spacer 2 (ITS2)-containing LSU pre-rRNAs move to PDFC-GC regions for processing. Inhibiting 5’ ETS processing disrupts the SSU pre-rRNA distribution and the nested FC-DFC sub-nucleolar structure. Cells in amniotes possess a multi-layered nucleolus, whereas anamniotes have only a bipartite structure with a merged FC-DFC. Kinetic labeling of pre-rRNA outflow shows a 7-fold higher pace in tetrapartite nucleoli over that in bipartite nucleoli, indicating that the emergence of the nested FC-DFC may facilitate an efficient SSU pre-rRNA processing over the course of evolution. Collectively, depicting the spatiotemporal distribution of pre-rRNAs reveals a key role of processing steps in organizing the multi-layered nucleolus and suggests a possible evolutionary advantage of the multi-layered structure in amniotes.

## Introduction

The mammalian nucleolus is spatially organized into four nested layers, namely, the fibrillar center (FC), the dense fibrillar component (DFC), the periphery of DFC (PDFC), and the granular component (GC)^1,2^. Each of these regions plays a distinct role in ribosome production^2,3^. In eukaryotic cells, 18S, 5.8S and 28S ribosomal RNAs (rRNA)s are transcribed together as a single precursor, known as 47S pre-rRNA in mammals, which is approximately 13 kb in length^4^. RNA polymerase I (Pol I) transcription occurs at the FC/DFC border^5^, and the transcribed 47S pre-rRNA moves outward across each layer within the nucleolus to form small (SSU)^6,7^ and large (LSU)^8^ ribosomal subunits in the GC^3,9^. The major precursor of SSU pre-rRNA is 30S, flanked by the 5’ external and internal transcribed spacers (ETS and ITS1), while the LSU precursor is mainly 32S pre-rRNA, producing 5.8S and 28S rRNAs (Extended Data Fig. 1a). Given that the 47S pre-rRNA is approximately 13kb long and would span roughly 4∼10 μm when fully extended^10^, it must fold considerably to fit within the much smaller nucleolus, of which the FC region measures around 150 nm and the FC-DFC region is roughly 400 nm in diameter^1,5^. This suggests that pre-rRNAs and their interacting proteins undergo remarkable compaction prior to SSU and LSU formation.

Specifically, nascent pre-rRNA is translocated to the DFC, where it undergoes chemical modifications and early cleavages at the 5’ and 3’ ends ^5,11^. It was thought that subsequent processing in the GC removes ETSs and ITSs, producing SSU and LSU^12,13^ (Fig. 1a and Extended Data Fig. 1a). However, biochemical studies are unlikely to unveil where the assembly of SSU and LSU occur in sub-nucleolus regions^4,12,13^, and the spatial distribution of pre-rRNAs within the nucleolus appears more complex^1,5^. The relationship between pre-rRNA processing dynamics and the multi-segmented nucleolar morphology has remained incompletely understood.

**Figure 1.**
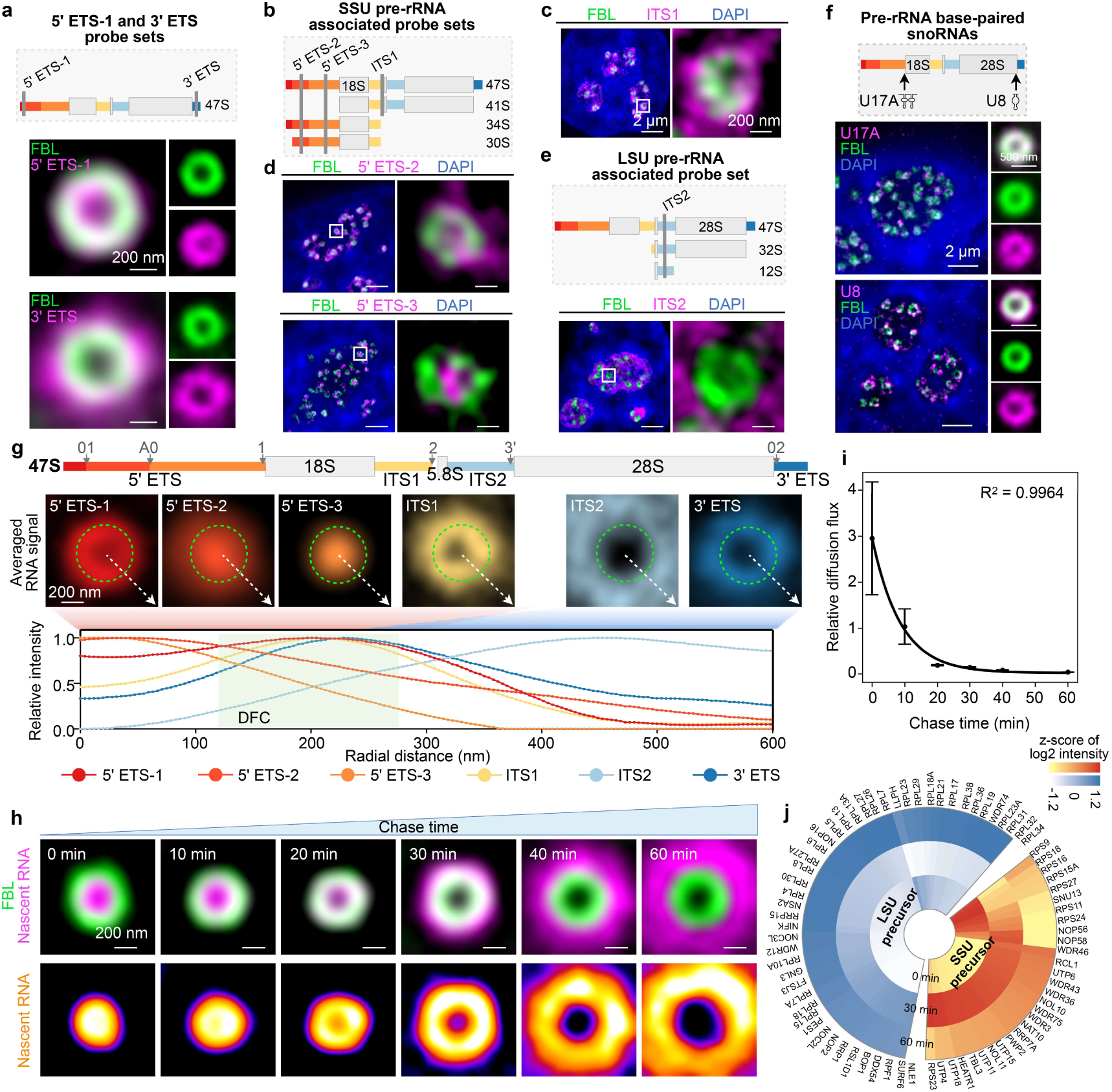
Spatiotemporal dissection of SSU and LSU pre-rRNA processing. **a**. The 5’ ETS-1 probe – and the 3’ ETS probe – detected 47S pre-rRNAs localize at the DFC and DFC/PDFC boundary, respectively. Top, schematic of 5’ ETS-1 and 3’ ETS smFISH probes. Bottom, representative smFISH images of 5’ ETS-1 and 3’ ETS in HeLa cells. FBL was used as the DFC marker. **b.** Schematic of the localization of smFISH probe sets targeting 5’ ETS-2, 5’ ETS-3 and ITS1. **c.** SSU pre-rRNAs processing predominantly occur in the FC to PDFC regions. The representative single-sliced SIM images of ITS1 (magenta) –detected SSU pre-rRNA is retained within the inner nucleolus ranging from the FC/DFC border to the PDFC in HeLa cells. **d.** Different 5’ ETS segments have distinct sub-nucleolar distributions, exemplified by smFISH images of 5’ ETS-2 and 5’ ETS-3. Of note, the 5’ ETS-2 extends from the FC to the PDFC, whereas the 5’ ETS-3 is confined to the FC. RNA, magenta; FBL, green. **e.** The ITS2 probe-detected LSU pre-rRNAs are localized to the outer nucleolar region ranging from the PDFC to the GC. Top, schematic of smFISH probe targeting ITS2. Bottom, representative single-sliced SIM images of ITS2 (magenta) and FBL (green) in HeLa cells. **f.** U17A and U8 snoRNAs targeting different pre-rRNA segments. Top, schematic depicting the targeted positions on the pre-rRNA of U17A and U8 snoRNAs. Middle and bottom, representative and averaged SIM images of U17A (middle, magenta) and FBL (green), U8 (bottom, magenta) and FBL (green) in HeLa cells. The number of DFC units (n) in the averaged images is 51 and 58, from 13 and 21 cells, respectively. **g.** Spatiotemporal distribution of pre-rRNA segments in different nucleolar sub-compartments. Top, averaged smFISH images of each pre-rRNA segment. Number of DFC units (n) = 33 (5’ ETS-1), 42 (5’ ETS-2), 55 (5’ ETS-3), 57 (ITS1), 52 (ITS2), 20 (3’ ETS), from 15, 15, 15, 21, 15, 11 cells, respectively. Bottom, the relative intensities of each segment plotted against radial position from the FC center, with the shaded green area indicating the DFC territory. The corresponding radial distance over the chasing time indicates nascent pre-rRNAs dynamics in Fig. 1h. The green circle marks the maximal intensity site of FBL. **h.** Representative average SIM images of nascent pre-rRNAs and FBL at different chasing time points. Nascent pre-rRNAs remain within the FC-DFC region during the first 30 min of labeling. Additional chasing to 40 and 60 min, pre-rRNAs reach the GC. The number of DFC units (n) analyzed across each time point is 58, 59, 59, 53, 52 and 50, from 17, 15, 13, 11, 13 and 13 cells, respectively. **i.** Decreased diffusion flux of pre-rRNAs across the nucleolus towards outside. The fitting curve was built based on the model shown in Extended Data Fig. 2d. Data are mean ± s.e.m. n = 20. **j.** Heatmap of pre-rRNA-bound proteins in dynamics (0-, 30-, and 60-min chase). Proteins associated with SSU precursors are largely enriched at 30 min, while those associated with LSU precursors are mostly enriched at 60 min. The complete heatmap of enriched proteins associated with SSU and LSU precursors are listed in Extended Data Fig. 3d. The log-transformed intensities were z-score transformed for the feature standardization, by the mean and standard deviation of peak enrichment time of all proteins.

Intriguingly, unlike the multi-layered mammalian cell nucleolus, anamniote cells, such as those in zebrafish, contain a single merged FC-DFC compartment named the fibrillar zone (FZ)^14,15^. The evolutionary expansion of the 5’ ETS and ITS2 regions^16^ indicates that an efficient processing system is likely required in the nucleolus during evolution. However, it is unclear whether the spatial processing of pre-rRNA in bipartite nucleoli differs from that in mammals, and whether such a processing is associated with an emergence of a more complicated nucleolus across species.

To understand whether and how the molecular scale rRNA processing is involved in the sub-nucleolar scale assembly of ribosomal subunits, we dissected the spatial processing relationship of pre-rRNA intermediates in the three-dimensional (3D) nucleolus. Unexpectedly, SSU and LSU pre-rRNAs are spatially segregated within FC-PDFC and PDFC-GC regions, with the SSU pre-rRNA processing essential for multi-layered nucleolus assembly. The spatial processing of pre-rRNAs in zebrafish embryos is much slower than that in human nucleoli, suggesting that spatial segmentation promotes more efficient pre-rRNA processing, which in turn likely supports organizing the multi-layered nucleolus.

## Results

### SSU pre-rRNA processing occurs in the FC-PDFC regions

To understand how the multi-step processing and transport of pre-rRNAs aligns with nucleolar organization, we designed multiple sets of smFISH probes to target specific pre-rRNA intermediates, and each probe was validated by Northern Blotting (NB) (Extended Data Fig. 1a-e). First were probes targeting 5’ ETS-1 and 3’ ETS regions that recognized 47S pre-rRNA (Fig. 1a and Extended Data Fig. 1b). Structured illumination microscopy (SIM) imaging of smFISH with individual probes revealed that the 5’ ETS-1-recognized pre-rRNA was predominantly localized within the DFC, while the 3’ ETS-targeted pre-rRNA was enriched near the DFC/PDFC boundary (Fig. 1a), confirming previous observations^1,5^ and also suggesting distinct pre-rRNA processing within the sub-nucleolar structure (Extended Data Fig. 1a).

Second, the cleavage within ITS1 at site 2 is crucial for separating the 18S rRNA (associated with SSU) from the 5.8S and 28S rRNAs (associated with LSU)^4^. Two sets of probes targeting ITS1 (Fig. 1b and Extended Data Fig. 1c, f) were designed to examine the localization of early pre-rRNA intermediates. These probe sets are capable of detecting 47S to 41S pre-rRNAs, confirmed by NB (Extended Data Fig. 1c). smFISH results showed that ITS1 signals were within and closely surrounding the DFC (Fig. 1c and Extended Data Fig. 1f). These observations suggested that the SSU processing preceded the release of LSU sequences into the next layer.

Next, to better understand the nucleolar distribution of different SSU-associated intermediates, such as 47S, 41S, 34S and 30S, we focused on the 5’ ETS, the major component of SSU pre-rRNA^17^. The 5’ ETS region can be divided into three segments according to its cleavage sites. 5’ ETS-1 (Fig. 1a) extends from the 5’-end to site 01, 5’ ETS-2 covers site 01 to A0, and 5’ ETS-3 spans from site A0 to 1^18^ (Extended Data Fig. 1a). Of note, 5’ ETS moves toward the DFC as it is transcribed ^5^ (Fig. 1a). With probes targeting 5’ ETS-2 and 5’ ETS-3, respectively (Fig. 1b), smFISH showed that 5’ ETS-2 signals extended from the FC to the PDFC, whereas 5’ ETS-3 signals were confined to the FC (Fig. 1d), which was unexpected from previous work. NB results confirmed that both probe sets mostly detected 47S and 30S pre-rRNAs, the major components of SSU pre-rRNAs (Extended Data Fig. 1d). This ruled out the possibility that the observed smFISH signals originated from the cleaved 5’ ETS fragments.

Together, the distribution patterns of the 5’ ETS detected signals (Fig. 1a, d) representing 47S and 30S pre-rRNAs (Extended Data Fig. 1b, d), as well as the ITS1 detected signals (Fig. 1c and Extended Data Fig. 1c, f) representing 47S and 41S pre-rRNAs, suggested that the 47S and 30S pre-rRNAs containing SSU processomes assembly^19^ mainly distributes from the FC to the PDFC.

### LSU pre-rRNA processing occurs in the PDFC-GC regions

Next, two probe sets targeting ITS2 were used to visualize LSU-associated pre-rRNA intermediates^4^ (Fig. 1e and Extended Data Fig. 1e, g). NB results confirmed that the ITS2 probe predominantly detected 32S and 12S pre-rRNAs, which are the main precursors of LSUs. Similarly, the site 3’ probe mainly detected 32S pre-rRNA (Extended Data Fig. 1e). Unlike SSU-associated pre-rRNAs, smFISH signals of both ITS2 and site 3’ probes were localized in the PDFC and the GC regions (Fig. 1e and Extended Data Fig. 1g), suggesting that LSU processing and assembly predominantly occurs in the outer nucleolar region.

Next, we visualized the sub-nucleolar localization patterns of several essential snoRNAs that base pair with different coding regions of pre-rRNAs during their processing^20^, to support the findings shown by different ETS and ITS probes mentioned above. U17A snoRNA mostly interacts with 18S rRNA region to facilitate its maturation in the 5’ end^21^. smFISH signals of U17A were enriched in the DFC (Fig. 1f). Additionally, U13 and E2 snoRNAs^22^, which interact with the 3’ end of 18S rRNA, were also localized to the DFC (Extended Data Fig. 1h, i). Such DFC localization patterns of these snoRNAs during 18S rRNA maturation substantiate the smFISH-based SSU pre-rRNA localization patterns (Fig. 1b-d). In contrast, U8 snoRNA, which assists in the processing of both the 5’– and 3’-ends of 5.8S/28S rRNA precursors^23^, was localized to the DFC/PDFC boundary, aligning with its processing role in the maturation of LSU precursors (Fig. 1e, f).

The spatial distribution of all key pre-rRNA intermediates within each sub-nucleolar domain is summarized in Fig. 1g.

### Spatiotemporal nucleolar segmentation of nascent pre-rRNAs

Next, we tracked the dynamics of pre-rRNA intermediates and their associated ribosomal subunits spatiotemporally (Extended Data Fig. 2a). Taking advantage of uridine analogs to label nascent transcripts^9,24–26^, this approach incorporated 5-ethynyl uridine (5-EU) into nascent RNAs. Given the rapid transcription and processing of pre-rRNAs^27^, we pulse-labeled cells for 10 minutes (min) to ensure homogeneous nascent pre-rRNA labeling, followed by 0-60 min chasing with 10 min intervals before fixation (Extended Data Fig. 2a). Fluorescent dye was conjugated to 5-EU via click chemistry to visualize the nascent pre-rRNAs within nucleolar sub-compartments using SIM (Fig. 1h).

In the pulse-chase labeling experiment with imaging, we observed that nascent pre-rRNAs progressively move outward from the transcription sites at the border of the FC and the DFC. Quantitative analysis of the intensity of the pre-rRNA signals along the radial axis of sub-nucleolar compartments showed a steadily outward flux of labeled RNA over time (Extended Data Fig. 2b). Further quantification of the absolute radius of nascent rRNA signals over time yielded a migration velocity of ∼1 Å/s across sub-compartments (Extended Data Fig. 2b, c), in agreement with a previous study^9^.

### Reduced diffusion flux of pre-rRNAs in the outer nucleolar region

As the nascent pre-rRNAs move outward to the GC, they are inevitably distributed in a progressively larger volume, resulting in a lower spatial density (Extended Data Fig. 2d). This lower density corresponded to the reduced radial diffusion flux in the outer regions within individual nucleoli, compared to the inner nucleolar regions. Thus, in addition to a constant linear flow speed of ∼1 Å/s, the relative diffusion flux of pre-rRNAs was indeed proportional to their spatial density (Fig. 1i and Extended Data Fig. 2d). In the inner and more crowded nucleolar regions, where the spatial volume is smaller, the diffusion flux is theoretically higher. In contrast, the larger outer regions have a relatively sparser pre-rRNA distribution, and thereby a lower diffusion flux (Fig. 1i).

This tendency was reflected as a faster outflux rate of pre-rRNAs in the inner nucleolar regions, compared to that of the outer regions. After 30 min of chasing, pre-rRNAs were ready to move out of the inner FC/DFC region, where SSU pre-rRNA processing occurs (Fig. 1g, h). In contrast, after a further 30 min chase, pre-rRNAs were just entering the outer GC regions, where LSU pre-rRNA processing occurs (Fig. 1g, h). Even after a 120 min chase, a significant portion of nascent pre-rRNAs still remained in the GC region rather than fully exiting into the nucleoplasm (data not shown).

### Spatiotemporal nucleolar segmentation of SSU and LSU complexes

At the same time, 5-EU-conjugated with biotin was used to precipitate proteins bound to nascent pre-rRNAs at 0 min, 30 min, 60 min, followed by mass spectrometry (MS) analysis (Extended Data Fig. 2a), revealing the dynamic assembly of SSU and LSU precursors and their associated proteins over the indicated pulse-chase timepoints as discussed below (Fig. 1j).

We first evaluated the dynamic localization of well-established nucleolar proteins in different sub-compartments^1^ at the indicated time points (0, 30, 60 min) by Western Blotting (WB) (Extended Data Fig. 3a, b). As expected, the FC localized proteins, primarily RNA Pol I transcription factors, bound to nascent pre-rRNAs at the 10 min chase stage (Extended Data Fig. 3a). Proteins in the DFC and PDFC are those involved in rRNA modification and processing, such as FBL and DDX21, and showed predominant binding to nascent pre-rRNAs at the 30 min chase point (Extended Data Fig. 3a, b). GC proteins, however, such as B23, bound to nascent pre-rRNAs after 30 min, highlighting the sequential progression of pre-rRNA processing across the nucleolar sub-compartments (Extended Data Fig. 3a, b).

Furthermore, recently published time-resolved RNA interactome data using pulse-chase metabolic labeling with 4-thio-uridine (4sU)^28^ also revealed a similar radial dispersion of nucleolar proteins. Consistent with the findings of this study, reported FC, DFC, and PDFC proteins^1^ bound to nascent pre-rRNAs sequentially within the first 40 min of chase, and nascent rRNA-bound GC proteins were present after the 40 min time window. This independent study further supported the spatiotemporal interaction of nucleolar proteins with nascent pre-rRNAs during their maturation (Extended Data Fig. 3c).

These observations (Extended Data Fig. 3a-c) prompted us to analyze the sub-nucleolar localization patterns of pre-rRNA intermediates (Fig. 1 h) and the assembly status of ribosome subunits over 60 min in greater detail (Fig. 1j and Extended Data Fig. 3d). Our analyses revealed distinct spatiotemporal distributions of different pre-rRNAs (Fig. 1h), corresponding to the segregated distribution pattern of SSU and LSU pre-rRNAs obtained from imaging data (Fig. 1g). These previously undescribed spatial distributions of SSU and LSU pre-rRNAs coupled with their *in situ* processing is beyond the current biochemical understanding of SSU and LSU assembly^7,8^ (Fig. 1g-j).

Specifically, the SSU-associated components mainly precipitated with nascent pre-rRNAs within 30 min (Fig. 1j and Extended Data Fig. 3d). During this stage, the nascent pre-rRNAs localized at the inner nucleolar regions ranging from the FC to the PDFC (Fig. 1h), covering a radius of approximately 250 nm (Fig. 1c, d, g and Extended Data Fig. 1f). This processing is rapid and detected as soon as the transcription occurs within the first 30 min chase (Fig. 1h), thus likely co-transcriptionally^5^. In contrast, the LSU-associated proteins were largely enriched at 60 min (Fig. 1j and Extended Data Fig. 3d) and processed more slowly in the outer nucleolar regions. At this time point, the nascent pre-rRNAs reached the outermost nucleolus at the GC (Fig. 1h), with a signal radius of approximately 450 nm, which corresponds to the spatial distribution pattern of LSU pre-rRNA (Fig. 1e, g and Extended Data Fig. 1g, 2b).

Additionally, our analyses also showed that SSU assembly and the processing of 18S rRNA precursors (Fig. 1a-d and Extended Data Fig. 1f, 3e) appeared to be more complex in distribution patterns compared to the LSU and 28S rRNA precursors in the nucleolus (Fig. 1e and Extended Data Fig. 1g, 3e).

### Inefficient 5’ ETS processing accompanies post-mitotic cell division

Given the observed spatial organization of pre-rRNA processing, we asked whether there exists any functional interdependency between processing and sub-nucleolar organization.

Human pluripotent stem cells (H9 cells) and differentiated arcuate neurons, represent two distinct cellular states: rapidly dividing stem cells and post-mitotic neurons with limited proliferation capacity^29^. In differentiated neurons, which have limited proliferation capacity, the nucleolus underwent a striking re-organization starting at the 2^nd^ day of differentiation (Fig. 2a and Extended Data Fig. 4a). In rapidly proliferating H9 and HeLa cells, the nucleoli were large and composed of several dozen FC/DFC units (Fig. 2a, b), whereas in differentiated neurons and SH-SY5Y cells, the nucleoli were dramatically reduced with only single digit numbers of FC/DFC units (Fig. 2a, b). Furthermore, differentiated neurons and SY-SH5Y cells contained larger individual FC volumes, compared to the highly proliferative H9 and HeLa cells (Fig. 2a-c and Extended Data Fig. 4b, c). Of note, the overall multi-layered nucleolar sub-structures persist throughout differentiation (Fig. 2a).

**Figure 2.**
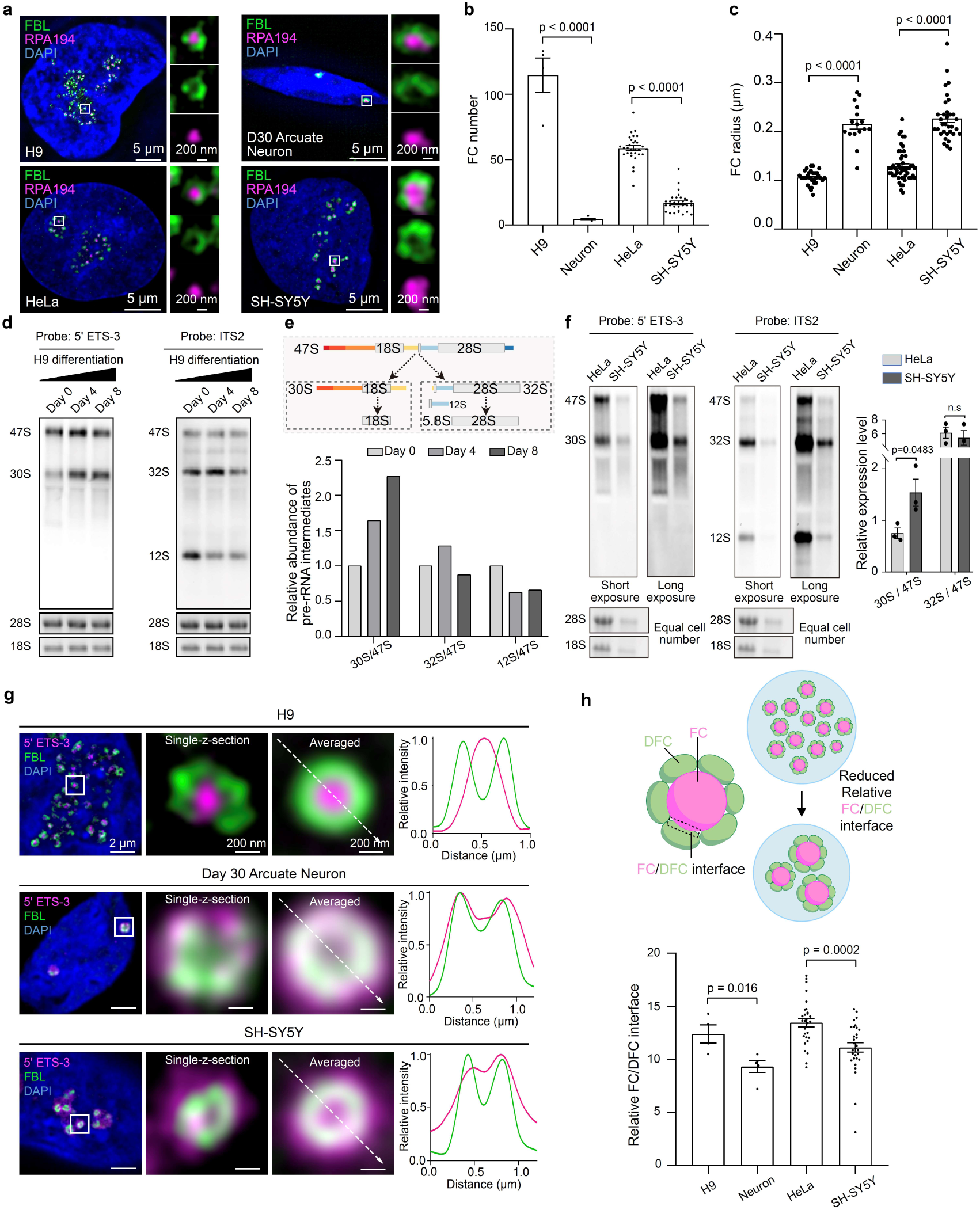
Inefficient SSU processing is associated with a decreased FC/DFC interface and altered 5’ ETS-3 localization. **a**. Representative SIM images of nucleoli in H9, differentiated neuron cells, HeLa cells and SH-SY5Y cells, marked by FBL (green) and RPA194 (magenta). **b**. The number of FCs in H9, H9-differentiated D30 neuron cells, HeLa and SH-SY5Y cells is shown. Data are mean ± s.e.m. (n) = 4, 5, 30 and 30, respectively. **c**. Statistics of FC radius in different cell states including H9, differentiated neuron cells, HeLa cells and SH-SY5Y cells. Data are mean ± s.e.m. Number of cells (n) = 5, 7, 5, 4. **d**. Inefficient SSU pre-rRNA processing upon H9 differentiation to neurons. NB of pre-rRNA intermediates in cells at the indicated differentiation points, using probes targeting 5’ ETS-3 and ITS2. Ethidium bromide (EB) staining of 28S and 18S rRNA is shown as the loading control. **e**. Quantification of pre-rRNA intermediates determined in panel (**d**). Normalized to Day 0. Top, schematic of the two pre-RNA classes recognizing SSU and LSU, respectively. **f**. The SSU pre-rRNA is accumulated in the neuroblastoma SH-SY5Y cells. NB of pre-rRNA intermediates in HeLa and SH-SY5Y cells, using probes targeting 5’ ETS-3 and ITS2. EB staining of 28S and 18S rRNA is shown as the loading control. Right, quantification of the relative 30S/47S and 32S/47S expression. Data are mean ± s.e.m. n = 3 experiments. **g**. Altered distribution of 5’ ETS-3-detected SSU pre-rRNAs from FC-dominant to DFC-PDFC-dominant in differentiated neurons and SH-SY5Y cells. Representative single-sliced and averaged SIM images of 5’ ETS-3 (magenta) and FBL (green) in H9, Day 30 arcuate Neuron and SH-SY5Y cells. In H9 cells, the 5’ ETS-3 signal is predominantly concentrated in the FC, while in neurons and SH-SY5Y cells, it is predominantly localized in the DFC-PDFC regions. The plot profile at the right-bottom of each panel shows the relative intensity of 5’ ETS-3 and FBL. Number of DFC units (n) = 51 (H9), 11 (neurons), 23 (SH-SY5Y) from 15, 8 and 10 cells, respectively. **h**. The relative FC/DFC interface is decreased in cells with inefficient SSU pre-rRNA processing. Left, a simplified model of cells with differently shaped nucleoli. Of note, the smaller size of FC/DFC units represent a larger relative FC/DFC interface (Extended Data Fig. 4d). Right, statistics of the relative FC/DFC interface in H9, neuron, HeLa and SH-SY5Y cells, showing a positive correlation between the FC/DFC interface and the cell proliferation rate. Data are mean ± s.e.m. Number of cells (n) = 4, 5, 30 and 30, respectively.

To determine if these proliferation-associated cell state changes were related to pre-rRNA processing, we performed NB using probes targeting the 5’ ETS and ITS2 regions (Fig. 2d). In differentiated neurons, we observed a 2-fold increase in 30S pre-rRNA, indicating inefficient 5’ ETS cleavage, a critical event during SSU pre-rRNA processing (Fig. 2d, e). In contrast, the levels of 47S pre-rRNA, as well as 32S and 12S pre-rRNAs (LSU) were decreased in these differentiated neuron cells (Fig. 2d, e). This overall reduction of all related pre-rRNAs likely reflects the decreased Pol I transcription in these post-mitotic cells (Fig. 2d).

Similar observations were also found in human neuroblastoma SH-SY5Y cells with low proliferation. Compared to the pre-rRNA processing observed in HeLa cells (Fig. 2f), SH-SY5Y cells showed a 2-fold increased 30S/47S ratio, indicating an increased accumulation of 30S pre-rRNA, while the level of 32S/47S, which reflects the proportion of LSU-associated pre-rRNAs, remained largely unchanged (Fig. 2f). These results confirmed that pre-rRNA processing varies between fast– and slow-proliferating cells, with an accumulation of 30S pre-rRNA due to an inefficient 5’ ETS processing in slowly proliferative cells (Fig. 2d-f).

### Inefficient 5’ ETS processing leads to altered SSU pre-rRNA distribution

The reduced 5’ ETS pre-rRNA processing (Fig. 2d-f) prompted us to ask whether the spatial distribution of pre-rRNA was also altered upon cell differentiation. The most noticeable change was the 5’ ETS-3 signals, which moved from the innermost FC region in stem cells (Fig. 2g) (and in HeLa cells, Fig. 1d) to the DFC-PDFC region in differentiated neurons and SH-SY5Y cells (Fig. 2g).

This altered localization of 5’ ETS-3 signals in neurons and SH-SY5Y cells (Fig. 2g) was correlated with the accumulation of 30S pre-rRNA detected by the same probe, showing increased 30S pre-rRNA levels and impaired 5’ ETS cleavage in these slowly proliferating cells (Fig. 2d-f). In H9 cells, smFISH signals of 5’ ETS-3 predominantly originated from 47S pre-rRNA (Fig. 2d, e, g), which is incorporated into the early-stage SSU processome (Extended Data Fig. 3e). However, neurons and SH-SY5Y cells exhibited a nearly doubled 30S/47S ratio compared to that in H9 and HeLa cells (Fig. 2d-f), showing that the 5’ ETS-3-detected 30S pre-rRNAs were enriched in the DFC in these cells (Fig. 2g). This observation indicated that the insufficient processing of 5’ ETS during SSU maturation may drive its outflux from the FC to the DFC-PDFC region, instead of staying within the FC-DFC region. In contrast, the localization of other pre-rRNA segments, including SSU-associated 5’ ETS-1 and ITS1 (Extended Data Fig. 5) and LSU-associated ITS2 and 3’ ETS (Extended Data Fig. 6) remained largely unchanged.

### Reduced FC/DFC interface in slowly proliferating cells

Along with the alteration of SSU pre-rRNA processing, the nucleolus underwent a striking re-organization. Given that 5’ ETS processing of pre-rRNAs occurs across the DFC (Fig. 1g) and is accompanied with an altered sub-nucleolar FC/DFC structure (Fig. 2a-c), we hypothesized that inefficient 5’ ETS-3 processing is accompanied by an altered FC/DFC organization. To test this idea, we analyzed the relative FC/DFC interface as one parameter to evaluate the change of individual FC/DFC substructures. Since SSU pre-rRNA processing mainly takes place at the FC-DFC interface (Fig. 1d), we reasoned that the contact area between the surfaces of FC and DFC in 3D might be an indicator of an altered FC/DFC substructure (Fig. 2h).

To minimize bias caused by variations in nucleolus size across different cell types, we normalized the FC/DFC contact area to the respective FC volume to generate a relative FC/DFC interface (Extended Data Fig. 4c). Theoretically, by simplifying the FC/DFC units into spherical structures, the relative FC/DFC interface was inversely proportional to individual FC radius (Extended Data Fig. 4c).

The FC and DFC regions were labeled with RPA194 (the largest subunit of Pol I) and FBL, respectively. Using the *Surface Surface Contact Area* plugin in Imaris, we experimentally quantified the relative FC/DFC interface in cells with varying proliferation rates (Fig. 2h). Our results showed that differentiated neurons with slow proliferation displayed a reduced relative FC/DFC interface (Fig. 2h). This experimentally validated relative FC/DFC interface in different cell types was inversely proportional to the FC radius, consistent with our prediction (Fig. 2c, h and Extended Data Fig. 4c).

These findings indicated a decreased FC/DFC spherical interface in slowly dividing cells, such as neurons, accompanied by inefficient 5’ ETS processing; as well as a connection between pre-rRNA processing and nucleolar substructure organization.

### 5’ ETS processing is important for the nested FC/DFC sub-structure

Given that nucleolar proteins alone are insufficient to organize the hierarchical nucleolus^30^, and that the FC/DFC interface is associated with the 5’ ETS processing efficiency (Fig. 2d-f, h), we proposed that SSU pre-rRNA processing may drive the assembly of FC/DFC.

To dissect which step of SSU pre-rRNA processing is key for the nested FC/DFC organization, we perturbed SSU pre-rRNA processing by designing ASO-Site A0 and ASO-Site 1 respectively targeting the key 5’ ETS cleavage sites (Fig. 3a). NB analyses confirmed significant increases in 30S and 26S rRNA precursors following ASO-Site A0 and ASO-Site 1 treatments, revealing successful perturbations as expected (Fig. 3b). The spatial distribution of 5’ ETS-centered SSU pre-rRNAs was dramatically changed after ASO treatment. While 5’ ETS-3-detected signals were still enriched in the FC in the scramble ASO-treated cells, smFISH signals became diffused to the outermost nucleolar regions upon ASO-Site A0 or ASO-Site 1 treatment (Fig. 3c). Of note, the 5’ ETS-3 probe-detected pre-rRNAs were shifted from 47S to 30S and 26S pre-rRNA intermediates upon the ASO-Site A0 and ASO-Site 1 treatments, respectively (Fig. 3b).

**Figure 3.**
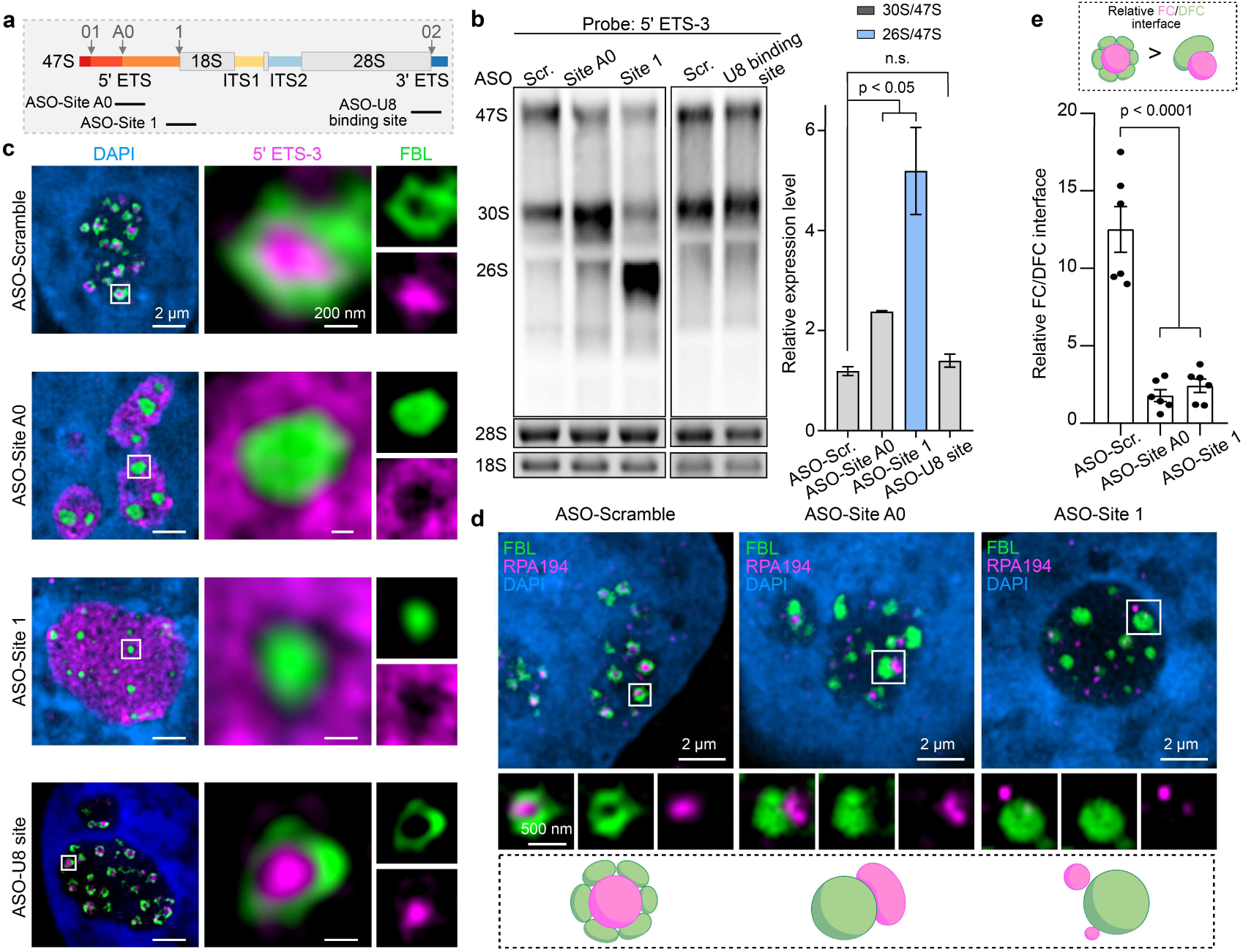
Impaired 5’ ETS processing alters SSU pre-rRNA composition and disrupts nested FC/DFC organization. **a,** Schematic of designed ASOs targeting A0 and 1 cleavage sites on the pre-rRNA. **b,** ASOs targeting the cleavage Site A0 and 1 of 5’ ETS impair the SSU pre-rRNA processing, unlike the ASO targeting LSU pre-rRNA region. Left, Northern Blot results of pre-rRNA intermediates, using probe recognizing 5’ ETS-3 are shown. Ethidium bromide staining of 28S and 18S rRNA is shown as the loading control. Right, quantification of pre-rRNA intermediates determined by NB. **c,** Impaired 5’ ETS processing via ASOs targeting the A0 and 1 cleavage sites results in reversed 5’ ETS-3 signals from the FC region “inside out”. Representative single-sliced SIM images of 5’ ETS-3 (magenta) and FBL (green) in HeLa cells treated with ASO-Site A0 and ASO-Site 1, as well as control treatments of the ASO-Scr. and the ASO-U8 site are shown. **d,** Impaired 5’ ETS processing disrupts the nested nucleolar structure. Representative single-sliced SIM images of RPA194 (magenta) and FBL (green) in HeLa cells treated with ASO-scramble (left), ASO-Site A0 (middle), and ASO-Site 1 (right). Bottom, the representative models of the FC-DFC units. **e,** Suppressed SSU pre-rRNA processing via ASOs results in a remarkable reduction of the relative FC/DFC interface in spherical 3D. Data are mean ± s.e.m. n = 6 cells.

Consistent with the LSU pre-rRNA localization patterns (Fig. 1g and Extended Data Fig. 7a), their localization (the ITS2 signal) was largely unaffected and remained in the GC (Extended Data Fig. 7a). Furthermore, a similar inhibition of 32S pre-rRNA processing by targeting the U8 binding site with an ASO showed no detectable changes on the spatial distribution of both 5’ ETS-3 and ITS2 detected signals (Fig. 3c and Extended Data Fig. 7a). These observations suggest that accurate 5’ ETS-related pre-rRNA processing is required for SSU pre-rRNA spatial distribution in the FC/DFC (Fig. 1g).

In agreement with the altered spatial distribution of SSU pre-rRNAs coupling with the disrupted nucleolar FC/DFC organization (Fig. 3c), further co-colocalization studies between the FC marker RPA194 and the DFC marker FBL showed an “inside-out” pattern with limited overlapping contacts between the two proteins at their interface (Fig. 3d). This observation supports the idea that SSU pre-rRNA processing plays a critical role in organizing the nested FC/DFC structure. Notably, such ASO-perturbed 5’ ETS processing also yielded a dramatic reduction of the relative FC/DFC spherical interface area (Fig. 3e), an increase in single FC volume and a decrease in FC numbers (Extended Data Fig. 7b, c). These analyses were consistent with reduced SSU pre-rRNA processing (Fig. 3b), and the nucleolar morphological changes observed in neuron cells (Fig 2a-c).

Similar analyses were also performed for LSU pre-rRNA processing, which occurs in the outermost nucleolar regions (Fig. 1e-g). Using ASO to target U8 snoRNA binding site that disrupts 28S pre-rRNA processing (Fig. 3b) showed no detectable morphological changes of nucleoli (Fig. 3c), suggesting that LSU pre-rRNA processing is dispensable for the nested FC/DFC structure.

### A bipartite nucleolus displays a distinct SSU pre-rRNA distribution pattern

Unlike multi-layered human nucleoli, anamniote cells, such as those in zebrafish, contain bipartite nucleoli with a single and large FC/DFC compartment named the fibrillar zone (FZ)^14,15^. Given that impaired 5’ ETS processing disrupted the FC/DFC organization (Fig. 3d) and reduced the relative FC/DFC interface area (Fig. 3e), we asked whether the spatial distribution of pre-rRNA segments in zebrafish embryos remained the same radial flux distribution as those in mammalian cells. In contrast to the pre-rRNA distribution in the human nucleolus (Fig. 1g), 5’ ETS signals in zebrafish did not localize within the inner region of Fbl, but rather partially overlapped and surrounded Fbl (Fig. 4a), while ITS2 signals were predominantly localized at the outer region of Fbl in the bipartite nucleolus (Fig. 4a), consistent with its distribution in HeLa cells (Fig. 1g). Additionally, compared to the ∼3.6 kb length of human 5’ ETS (Extended Data Fig. 1a) ^6^, zebrafish 5’ ETS is shorter, at approximately 1 kb^31^, which may also contribute to the simpler nucleolar structure in zebrafish.

**Figure 4.**
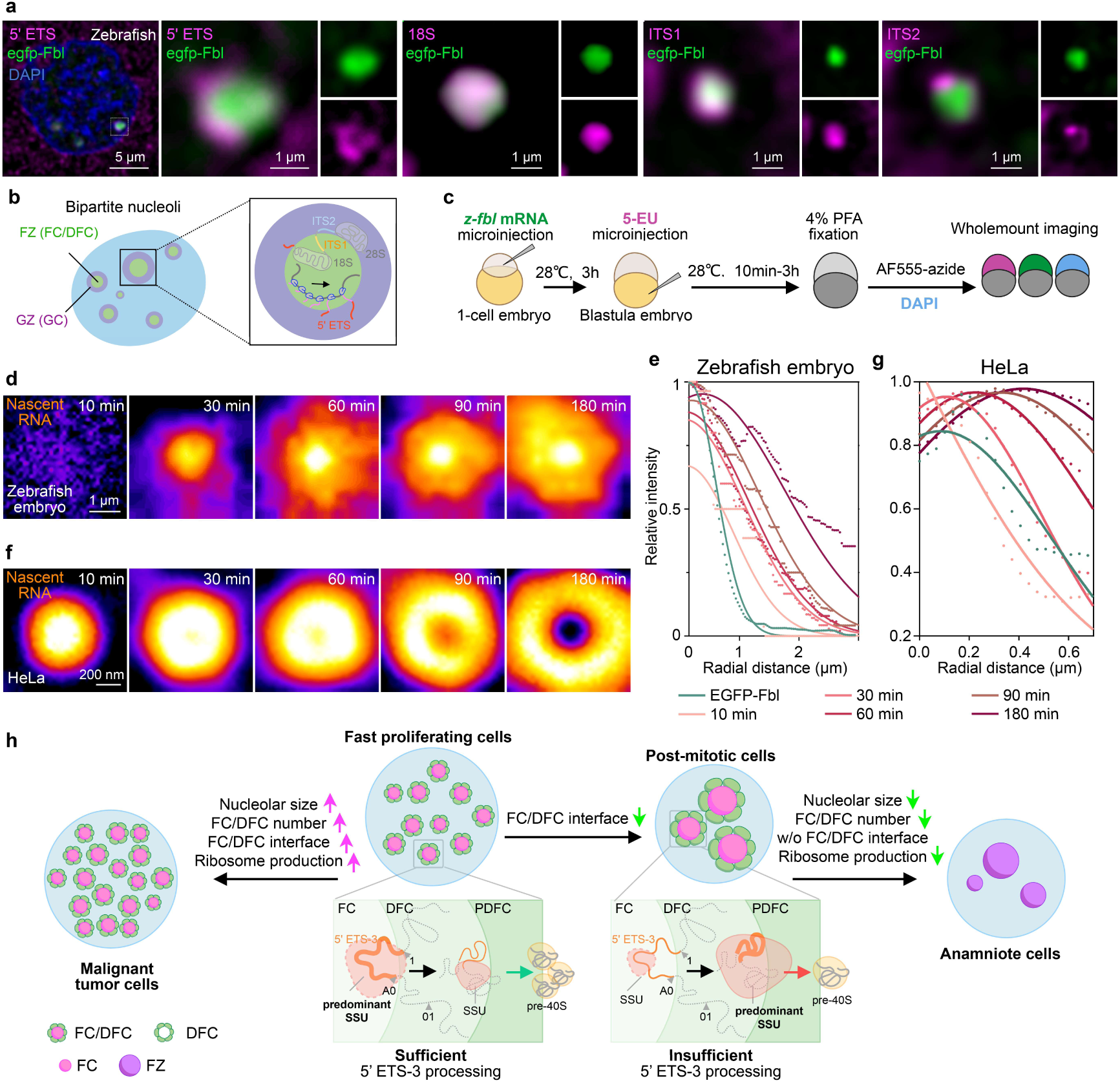
The multi-layered nucleolus ensures efficient SSU pre-rRNA processing over bipartite nucleolus. **a,** The spatial distribution of pre-rRNA in the bipartite nucleolus of cells from zebrafish embryos. smFISH images of different pre-rRNA segments in zebrafish embryos are shown. **b,** A proposed model of the bipartite nucleolus with a different spatial distribution pattern of pre-rRNA segments. **c,** The workflow of nascent pre-rRNAs labeling in cells of zebrafish embryos to examine the kinetics of nascent pre-rRNA processing. **d,** Representative averaged images of nascent pre-rRNAs at the indicated time points in the cell from zebrafish embryos are shown. The number of the FZ units (n) across each time points is 58, 59, 59, 53, 52, 50, respectively. **e,** Quantification of pre-rRNAs intensities from images in panel (**d**), with the radial position from the Fbl center to the edge of detected signals at each indicated labeling time. **f,** Representative averaged images of nascent pre-rRNAs at the indicated labeling time points in HeLa cells. The number of FC/DFC units (n) across each time points is 34, 32, 36, 33, 32, respectively. **g,** Quantification of pre-rRNAs intensities from images in panel (**f**) with radial position from the FC center to the edge of detected signals at each indicated labeling time. **h,** A proposed model of the coordination of pre-rRNA processing, spatial distribution and nucleolar sub-structures. In fast proliferating cells, efficient 5’ ETS processing of SSU pre-rRNA occurs to meet the demand of ribosome synthesis, with 5’ ETS-containing pre-rRNAs spatially distributed from the FC to the PDFC. This corresponds to the enlarged FC/DFC interface. In slow proliferating cells, the 5’ ETS processing becomes inefficient, accompanying with the reorganized FC/DFC interface. Consistently, a bipartite nucleolus in anamniote cells completely lacks the FC/DFC interface, showing a centered distribution of SSU pre-rRNA radial flux and the delayed processing upon transcription. In (e) and (g), the curves are fitted with Gaussian non-linear regression. n = 15.

Based on the bipartite nucleolar organization^14,15^ and distinct spatial distributions of pre-rRNA segments (Fig. 4a), we proposed that zebrafish SSU pre-rRNA spatial distribution might be markedly different from that in the human nucleolus (Fig. 4b). The nascent pre-rRNA is transcribed within the FZ, and the 5’ ETS region co-transcriptionally moves out to the periphery of the FZ. This distinct nucleolar structure, with its merged FC-DFC (FZ) in zebrafish (Fig. 4a, b), likely reflects a less efficient SSU pre-rRNA processing compared to the highly organized FC/DFC interface seen in mammalian nucleoli (Fig. 1g).

### The multi-layered nucleolus outperforms the bipartite one in pre-rRNA processing

We next performed 5-EU labeling to quantify the outflux rate of pre-rRNAs from the inner to the outer nucleolar sub-compartments within the bipartite nucleoli. EGFP-Fbl mRNA was microinjected into 1-cell stage zebrafish embryos to label the FZ. At approximately 3 hr post-fertilization (hpf), when zygotic genome activation (ZGA) began, we injected 5-EU into the same embryos to label newly transcribed pre-rRNAs. Embryos were then fixed at various time points to label nascent pre-rRNAs (Fig. 4c).

In zebrafish, nascent pre-rRNAs accumulate within the Fbl-marked FZ after 30 min of labeling. Over 60-90 min, a fraction of transcribed rRNAs gradually moved across the FZ (∼1 μm border) into the GZ, without exceeding the 3 μm boundary (Fig. 4d, e and Extended Data Fig. 8a). Even after 180 min of tracing, the nascent pre-rRNA signals still remained restricted to the FZ, with a significant portion persisting in the transcriptional region (Fig. 4d, e and Extended Data Fig. 8a). To compare the pre-rRNA flux rate between the two distinct nucleoli, we performed the same labeling assay in HeLa cells (Fig. 4f, g and Extended Data Fig. 8b). Two remarkable differences between these two evolutionarily different nucleoli were observed. First, the nascent pre-rRNAs in the bipartite nucleoli showed a noticeable accumulation in the innermost transcriptional nucleolar regions, even with the extended 5-EU labeling for 60–180 min. In contrast, pre-rRNAs were consistently moved to the outer nucleolar regions in the multi-layered nucleolus (Fig. 4d-g).

Second, the outflux rate of pre-rRNAs in the bipartite nucleolus of zebrafish was notably delayed. In zebrafish nucleoli, labeled nascent pre-rRNAs remained in the inner nucleolar regions where transcription occurs (Fig. 4d, e). However, nascent pre-rRNAs in HeLa cells efficiently moved away from the transcription site at the FC border to the GC (Fig. 4f, g). Quantitation of these imaging results revealed the quick radial flux movement of the maximal peak intensity along the labeling time progression in HeLa cells (Fig. 4g), compared to that of zebrafish nucleoli (Fig. 4e). Quantification of the maximal peak intensity at each labeling point in zebrafish and HeLa cells (see Methods) revealed a 7-fold reduction of the pre-rRNA outflux rate in bipartite nucleolus compared to that in the multi-layered nucleolus (Fig. 4e, g). Of note, the intensity of the Pol II transcribed nascent RNAs were kept increasing over the 180 min continuous 5-EU labeling (Extended Data Fig. 8c), excluding a possible 5-EU exhaustion that might cause the signal reduction of nascent pre-rRNAs within the innermost FC region in HeLa cells (Fig. 4f and Extended Data Fig. 8b).

Collectively, these results support the view that the multi-layered nucleolus facilitates efficient rRNA processing and that an efficient rRNA processing likely substantiates such a multi-layered nucleolar organization.

## Discussion

The nucleolus is one of the most well-studied nuclear bodies, yet the mechanism underlying its multi-layered organization and the role of such a complex structure in supporting efficient pre-rRNA processing have remained elusive. This gap in knowledge has largely stemmed from a lack of capable toolkits to probe the *in-situ* distribution-processing relationships at play. The combination of high-resolution pre-rRNA intermediates imaging (Fig. 1a-g and Extended Data Fig. 1f-i), the spatiotemporal mapping of pre-rRNA processing (Fig. 1h-j and Extended Data Fig. 2, 3) and the mathematical modeling of the FC/DFC interface (Fig. 2h and Extended Data Fig. 4c) have unveiled an asynchronous detail of the pre-rRNA molecular-scale processing within the micron-scale sub-nucleolar structure (Fig. 1g-j, Fig. 4h).

Our results suggest that SSU pre-rRNA processing in the inner sub-compartments (Fig. 1b-d) plays a key role in determining its spatial distribution (Fig. 2g, 3c) and organizing the nested FC/DFC structure (Fig. 2c, h, 3d). An efficient SSU pre-rRNA processing in the nested FC/DFC is required for cells with fast proliferation (Fig. 2a-f), and likely the evolutionarily emerged multi-layered nucleolus (Fig. 4). These observations suggest an unexpected functional interdependency between SSU pre-rRNA processing and the existence of nested FC/DFC units with adjustable shapes (Fig. 4h). However, these data cannot distinguish RNA synthesis and decay, as both contribute to the steady-state levels of intermediates. Further studies are warranted to dissect these possibilities.

The finding that pre-rRNA intermediates and their associated ribosomal proteins accumulate at distinct sub-nucleolar domains (Fig. 1g-j) shows an underappreciated spatial-structure relationship for nucleolar function and organization^12^. The assembly of the SSU mainly occurs from the FC to the PDFC (Fig. 1b-d, g), while LSU assembly largely takes place at the PDFC to the GC (Fig. 1e, g). This separation of SSU and LSU pre-rRNA processing reflects the precise regulation of rRNA maturation in space and time within the crowded nucleolar architecture for ribosome biogenesis. Importantly, the SSU pre-rRNA processing, particularly the cleavage of the 5’ ETS region, is essential for maintaining the nested FC/DFC organization with enhanced FC/DFC interface, which in turn supports the rapid SSU pre-rRNA processing (Fig. 2g, h). Similar observations were also very recently reported^32^, in which RNA sequencing was applied to measure the kinetics of nascent pre-rRNA cleavage and modification within the nucleolus.

Such an efficient SSU pre-rRNA processing and the nested FC/DFC organization is highly adaptable with cell proliferation. In slowly proliferating cells such as neurons, impaired 5’ ETS processing correlates to 30S pre-rRNA accumulation (Fig. 2d-f) and reduced proliferative capacity (Fig. 2a-c, h, 3d-e), highlighting an interdependence between the SSU processing and the nucleolar substructure in response to the cellular demand. From an evolutionary point of view, such an efficient and highly organized SSU processing appears to necessitate the emergence of the multi-layered nucleolus. Cells from zebrafish embryos possess a less efficient SSU processing compared to those from amniotes (Fig.4d-g). Thus, the presence of distinct FC-DFC subdomains with increased FC/DFC interface in amniotes may provide an advantage by enhancing SSU pre-rRNA processing for the emergence of the complex, multi-layered nucleolus (Fig. 4h). Future studies are warranted to better understand how the appearance of the multi-layered nucleolus would impact the ribosome biogenesis necessary for amniotic cellular requirement.

## Acknowledgment

We thank members of the Chen laboratory for discussions; Sarah A. Woodson, Yi-Lan Chen, and Gordon Carmichael for the critical reading of our manuscript; Narry Kim for kindly providing the data of nascent RNAs bound proteins; L.-Z. Yang, M. L. Hou and B.-Y. Zou for supporting our experiments. This work was supported by the Strategic Priority Research Program of the Chinese Academy of Science (XDB0570000), the National Key R&D Program of China (2021YFA1100203), the Postdoctoral Innovation Talent Support Program, and the Shanghai “Super Postdoc” Incentive Program. This work was also supported by the New Cornerstone Science Foundation through the New Cornerstone Investigator Program and the XPLORER PRIZE.

## Author contributions

L.-L.C. supervised and conceived the project. L.-L.C., Y.-H.P. and L.S. designed experiments. Y.-Y.Z. performed computational analyses supervised by L.Y. Y.-H.P., L.S., performed all other experiments and analyses with the help of Z.-H.Y., Y.Z., S.-M.C. and J.Z. L.-L.C., Y.-H.P. and L.S. drafted the manuscript. L.-L.C. edited the manuscript.

## Methods

### Cell culture

Human HeLa, SH-SY5Y and HEK293FT cell were purchased from the American Type Culture Collection (ATCC; http://www.atcc.org) and were authenticated using STR profiling. Human H9 cells were purchased from WiCell. HeLa and HEK293FT cells were maintained in DMEM, which was supplemented with 10% fetal bovine serum (FBS). H9 cells were maintained in DMEM/F-12 supplemented with 20% KnockOut Serum Replacement, 0.1 mM Glutamax, 0.1 mM non-essential amino acids and 0.1 mM mercaptoethanol and 4 ng/ml b-FGF and cultured with irradiated mouse embryonic fibroblast feeder cells with daily changed cultured medium and passaged weekly. SH-SY5Y cells were maintained in a 1:1 mixture of Eagle’s Minimum Essential Medium and F12 Medium with 10% FBS. All cells were cultured at 37 °C in a 5% CO_2_ cell culture incubator and were routinely tested to exclude mycoplasma contamination.

### Differentiation of H9 cells

Cell differentiation was performed as described before^33^. H9 cells grown on 6 well coated with matrigel. The neuron differentiation procedure started when H9 cells reached 95% to 100% confluence. H9 cells were cultured in KSR medium with 100 ng/ml SHH (R&D Systems; cat. no. 1845-SH), 2 μM purmorphamine (Selleckchem; cat. no. S3042), 10 μM SB431542 (Selleckchem; cat. no. S1067), and 2.5 μM LDN-193189 (Selleckchem; cat. no. S2618) and the medium changed daily from day 1 to day 4. The medium was changed to KSR medium and N2 medium from 3:1, 1:1 to 1:3 ratio and supplemented with 100 ng/ml SHH, 2 μM purmorphamine, 10 μM SB431542, and 2.5 μM LDN-193189 from day 5 to day 7. On day 8, the medium was changed to N2 medium supplemented with 100 ng/ml SHH, 2 μM purmorphamine, 10 μM SB431542, and 2.5 μM LDN-193189. From day 9 to day 12, the cells were cultured in N2 medium supplemented with 1× B-27 and 10 μM DAPT. The medium needed to be replaced daily. On day 10, we coated 0.01% poly-L-ornithine on new 10 cm dishes and the next day, aspirated poly-L-ornithine, washed the plates with sterile distilled water four times, added matrigel to each well, and incubated the plates overnight at 37°C. On day 12, the differentiated cells were treated with accutase for 8 min at 37°C, then added 2 ml N2 medium and detached all cells by repeatedly pipetting. After pelleting cells by centrifugation for 3 min at 800 rpm and aspirating the supernatant, N2 medium supplemented with B-27 and 10 μM ROCK inhibitor were used to suspend cell pallets. 6 million cells were seeded into every 10 cm dish coated with matrigel. From day 13 to day 16, the N2 medium supplemented with B-27 and 10 μM DAPT was replaced daily. From day 17, the medium was changed to N2 medium supplemented with B-27 and 20 ng/ml BDNF every two days until fully differentiated neurons were observed at day 30.

### RNA isolation, Northern blotting (NB)

Total RNAs from the cultured cells were extracted with TRizol Reagent (Invitrogen) according to the manufacturer’s protocol. NB was carried out according to the manufacturer’s protocol (DIG Northern Starter Kit, Roche) to examine pre-rRNAs. RNA was loaded on agarose gels and the Dig-labeled antisense probes were used. Probe sequences for northern blot are listed in Supplementary Table 1.

### Western blotting (WB)

Nascent RNA-bound proteins were collected after treatments and resuspended in lysis buffer (1% NP-40, 0.5% sodium deoxycholate, 0.1% SDS, 150 mM NaCl, 50 mM Tris and 1× protease inhibitor cocktail, pH 8.0) for 10 min. After centrifugation, supernatants containing soluble proteins were resolved on polyacrylamide gel with 10% SDS and analyzed by WB with anti-FBL (Abcam, 1:1,000 dilution), anti-DDX21 (ProteinTech, 1:1,000 dilution), anti-B23 (Santa Cruz, 1:1,00 dilution) or anti-ATCB (Sigma, 1:5,000 dilution) antibodies.

### Single molecule RNA Fluorescence *in situ* Hybridization (smFISH)

All smFISH probes were designed via Stellaris Probe Designer and labeled with Cy3 on the 3’ end. smFISH was carried out as described before ^34^. In brief, cells were fixed with 4% PFA for 15 min, followed by permeabilization with 0.5% Triton X-100 for 10 min. Cells were incubated in 10% formamide/2× SSC for 10 min at room temperature followed by hybridization at 37 °C for 16 h. After hybridization, the cells were washed for 2 times with 10% formamide/2× SSC at 37 °C, each time for 30 min. Samples were mounted in VECTASHIELD antifade mounting medium (Vector Lab). Probe sequences for northern blot are listed in Supplementary Table1.

### Protein visualization

To detect protein localization by immunofluorescence in fixed cells, cells were seeded on High Performance No.1.5 18×18 mm glass coverslips and were fixed with 4% PFA for 15 min, followed by permeabilization with 0.5% Triton X-100 for 10 min. Then cells were blocked with 1% BSA for 1 hour at room temperature (RT). Primary antibodies were diluted with 1% BSA (FBL, Abcam, 1:300 dilution; RPA194, Santa Cruz, 1:50 dilution) and incubate for 1 hour at RT. After washing with 1× DPBS 3 times, fluorescent secondary antibodies were 1: 1000 diluted in 1% BSA and incubated for 1 hour at RT. Samples were mounted in VECTASHIELD antifade mounting medium (Vector Lab).

### Structured Illumination Microscopy (SIM) procedure

All SIM experiments were performed using commercialized Hessian-SIM, termed His-SIM (High Intelligent and Sensitive Microscope) equipped with a 100×/1.5NA oil immersion objective (Olympus). Image acquisition was carried out using IMAGER software. SIM image stacks were captured with a z-distance of 0.1 μm or 0.2 μm with 5 phases, 3 angles, and 15 raw images per plane.

### Spinning disk confocal super-resolution microscopy

Spinning disk confocal super-resolution microscope images were acquired with the Olympus IXplore SpinSR microscope equipped with Yokogawa CSU-W1 SoRa (50 μm and SoRa disks) and operated with cellSens Dmension Software.

### Pulse-chase labeling combined with imaging

HeLa cells were cultured in the medium supplemented with 1 mM 5-EU (Beyotime) for 10 min. Then the medium was replaced with the fresh 5-EU-free medium. Next, cells were fixed at indicated time points with 4% PFA for 15 min, followed by permeabilization with 0.5% Triton X-100 for 10 min. The click reaction for nascent RNA labeling was carried out as described in the manufacturer’s protocol (Thermo Fisher C10642) with the following modifications: (1) the free copper (component C) and the copper protectant (component D) level in the reaction mix was medium (1:1); (2) the Alexa Fluor picolyl azide concentration was 1 μM; (3) cells were incubated with the reaction mixture for 30 min and was washed 3 times with DPBS. Finally, samples were mounted in VECTASHIELD antifade mounting medium (Vector Lab) for imaging.

### Pulse-chase labeling combined with mass spectrometry (MS) analysis

The procedure was modified from previous publication ^35^. HeLa cells were seeded on 10-cm dishes to perform pulse-chase labeling until the confluence reaches 70-80%. Cells were pulse labeled with 5-EU (Beyotime) for 10 min at 37 °C with 5% CO_2_. Then, cells were washed twice with 1× DPBS to remove the remnant 5-EU. Cells were then chased with 5-EU-free medium over different time points. Next, cells were crosslinked with 1% formaldehyde for 10 min at room temperature, followed by 125 mM glycine treatment to stop the crosslinking reaction. Cells were washed twice with 1× DPBS, and then permeabilized with 0.5 % Triton X-100 for 10 min at room temperature, followed by twice wash with 1× DPBS. The click reaction mix was prepared with 20 mM Tris-HCl (pH 7.5), 250 μM biotin picolyl azide (Sigma-Aldrich, 900912), 1 mM CuSO_4_ (Sigma-Aldrich, 451657), 2 mM BTTAA (Vector Laboratories, CCT-1236), 1 mM aminoguanidine (Sigma-Aldrich, 396494), and 2.5 mM Sodium L-ascorbate (Sigma-Aldrich, 11140). 5-EU labeled nascent RNAs reacted with the mixture to conjugate biotin for 60 min at room temperature. Then the unbound biotin was washed out with 1× DPBS.

Cells were scrapped, resuspended in 1 mL lysis buffer (20 mM Tris pH 7.5, 500 mM NaCl, 1 mM EDTA pH 8.0, 0.5 mM PMSF, 2 mM RVC, protease inhibitor cocktail (Roche)) followed by 5× 20s sonication with an interval of 1 min on ice. The supernatant was collected after centrifuging at 14000 rpm for 10 min at 4 °C. 100 μL

Dynabeads MyOne Streptavidin C1 beads (Invitrogen, 65002) were washed with 1 mL lysis buffer for three times, followed by blocking beads with 1% BSA and 20 μg/ml yeast tRNA for 1 h at 4 °C. Then the supernatant was incubated with the beads for 3 h at 4 °C, followed by washing with wash buffer 1 (20 mM Tris pH 7.5, 500 mM NaCl, 1 mM EDTA pH 8.0, 0.5% SDS, 0.5 mM PMSF, 2 mM RVC, protease inhibitor cocktail (Roche)), wash buffer 2 (20 mM Tris pH 7.5, 500 mM NaCl, 1 mM EDTA pH 8.0, 0.1% SDS, 0.5 mM PMSF, 2 mM RVC, protease inhibitor cocktail (Roche)), and wash buffer 3 (20 mM Tris pH 7.5, 200 mM NaCl, 1 mM EDTA pH 8.0, 5 mM DTT, 0.5 mM PMSF, 2 mM RVC, protease inhibitor cocktail (Roche)). Finally, the enriched protein samples were sent to Shanghai Applied Protein Technology for mass spectrometry analysis.

### MS data analysis

MS raw data was analyzed with Maxquant 1.6.14. The main parameters were set as follow: the used database was from UniProt, enzyme specificity was set to ‘Trypsin’, allowing up to two missed cleavages; dynamic modifications were set at oxidation (M), fixed modifications were set at carbamidomethyl (C). Label-free quantification was enabled and ‘Match between runs,’ ‘iBAQ’ options were selected. The ProteomicsTools is 3.1.6. The filter by score was ≥ 20.

Protein information was obtained from “ProteinGroups.txt”. For protein groups containing multiple matched proteins, the LFQ intensity of the group was used to represent the best matched protein. For each protein, we first performed log2 transformation on LFQ intensity, then we applied z-score standardization at protein-wise direction using mean and standard deviation of protein intensity at all time points. We used the nucleolar protein sub-nucleolar localization dataset, SSU-associated protein dataset and LSU-associated protein dataset as three curated datasets. By comparing with these databases, 90 proteins with precise sub-nucleolar localization, 55 SSU-associated proteins and 60 LSU-associated proteins were found in our MS data. For each nucleolar sub-compartment localized proteins, LOESS regression (span = 0.75) of z-score of log2 protein intensity against time was performed in R (v4.3). For SSU precursor proteins and LSU precursor proteins, the time point when the z-score reached its maximum value was defined as peak enrichment time of the corresponding protein.

### ASOs treatment to inhibit pre-rRNA processing

For ASOs treatment, HeLa cells were seeded in 6-well plates, then the modified ASOs (3 μg/well) was transfected with Lipofectamine RNAiMAX Transfection Reagent (Invitrogen, 13778075) for 24 h before the subsequent assay.

### smFISH in zebrafish embryos

Zebrafish embryos were injected EGFP-Fbl mRNA (400 ng/μL) at 1-cell stage to label the FZ region of zebrafish nucleoli, followed by 4 h incubation at 28 °C. Embryos were fixed with 4% PFA for 4 h at room temperature. Embryos were dehydrated with methanol and kept at –20 °C for over 2 h. After rehydration, the embryos were incubated in prehybridization buffer (10% formamide, 2×SSC, 0.1% Triton X-100) at 30 °C for 5min. Then the embryos were soaked in probe solution and incubated at 30 °C overnight in dark. On the following day, embryos were washed twice with prehybridization buffer at 30 °C for 30 min each and then briefly washed with PBST. Embryos were mounted with 1% agarose with DAPI and photographed using confocal microscopy (Olympus SpinSR)

### Pulse labeling nascent RNAs with 5-EU in zebrafish embryos and HeLa cells

For pulse labeling nascent RNAs with 5-Eu in zebrafish embryos, we first injected EGFP-Fbl mRNA (400 ng/μL) into the embryos at 1-cell stage to label the FZ region of zebrafish nucleoli. At approximately 3 h incubation at 28 °C, when zygotic genome activation (ZGA) begins, we injected 5-EU (50 mM) into the yolk sac of zebrafish embryos to label nascent RNAs. Embryos were then fixed at 10 min, 30 min, 60 min, 90 min and 180 min post 5-EU injection with 4% PFA for 4 h at room temperature. Embryos were dehydrated with methanol and kept at –20 °C for over 2 h. after rehydration, the embryos were incubated with the click reaction mix (same as the mix in pulse-chase labeling combined with imaging in HeLa cells) for 30 in at room temperature. Subsequently, embryos were rinsed two times with PBST and photographed in 1% agarose gel using confocal microscopy (Olympus SpinSR).

For pulse labeling nascent RNAs with 5-EU in HeLa cells, HeLa cells were cultured in the medium supplemented with 1 mM 5-EU for 10min, 30 min, 60 min, 90 min and 180 min. Next, cells were fixed with 4% PFA for 15 min at room temperature, followed by permeabilization with 0.5% Triton X-100 for 10 min. The subsequent processes are the same with the protocol of pulse-chase labeling combined with imaging.

### Fitting of Diffusion Flux Data

The radial position of the peak intensity of nascent pre-rRNA in each nucleolus, starting from the FC to the edge of the GC, was obtained via the plot profile analysis by ImageJ. The time-distance dataset was used to calculate the flow velocity and to fit the diffusion flux curve.

The temporal dynamics of relative diffusion flux were modeled using an exponential decay function. According to the simplified pre-rRNA flow model across nucleolar sub-compartments, the fitting process assumed that the relative diffusion flux *J(t)* at time *t* follows the equation:

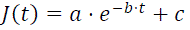

where:

- *J(t)* represents the relative diffusion flux at time *t*,
- *a* is the amplitude of the initial flux,
- *b* is the rate constant that governs the exponential decay,
- *c* is the steady-state flux, representing the asymptotic value that the flux approaches as *t* increases.

The fitting was performed using the nonlinear least squares method through the *scipy.optimize.curve_fit* function in Python, which optimizes the parameters *a*, *b*, and *c* to minimize the residuals between the model and the experimental data.

### Calculation of the relative FC/DFC spherical interface

The FC and DFC were labeled by the well-established marker proteins RPA194 and FBL, respectively. The interface was calculated with the XTension *Surface Surface Contact Area* by Imaris. The primary spherical surface is constructed according to the RPA194 signal, and the secondary spherical surface is constructed according to the FBL signal that covers the primary spherical surface. The final relative FC/DFC spherical interface was calculated by normalizing the total contact interface area in a cell to the total FC volume.

The average FC radius in each cell was calculated according to the total FC volume and the FC number by simplifying the FC as an ideal sphere. Of note, we simplified the FC-DFC as a nested spherical structure, the theoretical FC/DFC interface was the surface area of FC, and the theoretically relative FC/DFC interface was normalized to the radius of the FC.

### Fitting the curve of relative RNA intensity along the radial distance in pulse labeling assay in zebrafish and HeLa cells

The relative intensity of nascent pre-rRNAs along the radial distance was obtained by ImageJ. Data from 14 zebrafish embryos at each time point were plotted. Data from 15 FC/DFC units of more than 5 HeLa cells at each time point were plotted. The fitted relative intensity curve is obtained by Gaussian non-linear regression.

The movement rate of nascent pre-rRNAs in zebrafish embryo and HeLa cells was calculated by examining the radial position of maximal peak intensity at the sequential labeling time points and normalizing the radial position to the radius of FBL. The normalized radial distance at each time point was used to calculate the movement rate.

### Statistics

Statistical analyses were performed with GraphPad Prism 8, R (v4.3) and Python 3.8.8. For the sample size, statistical method and significance of all graphs, please see figure legends and methods for details. We used Student’s t-test to analyze between-group differences. *P*-value < 0.05 was considered to indicate statistical significance. No data was excluded from the analysis.

## Extended Data Figures and Figure legends

**Extended Data Figure 1.**
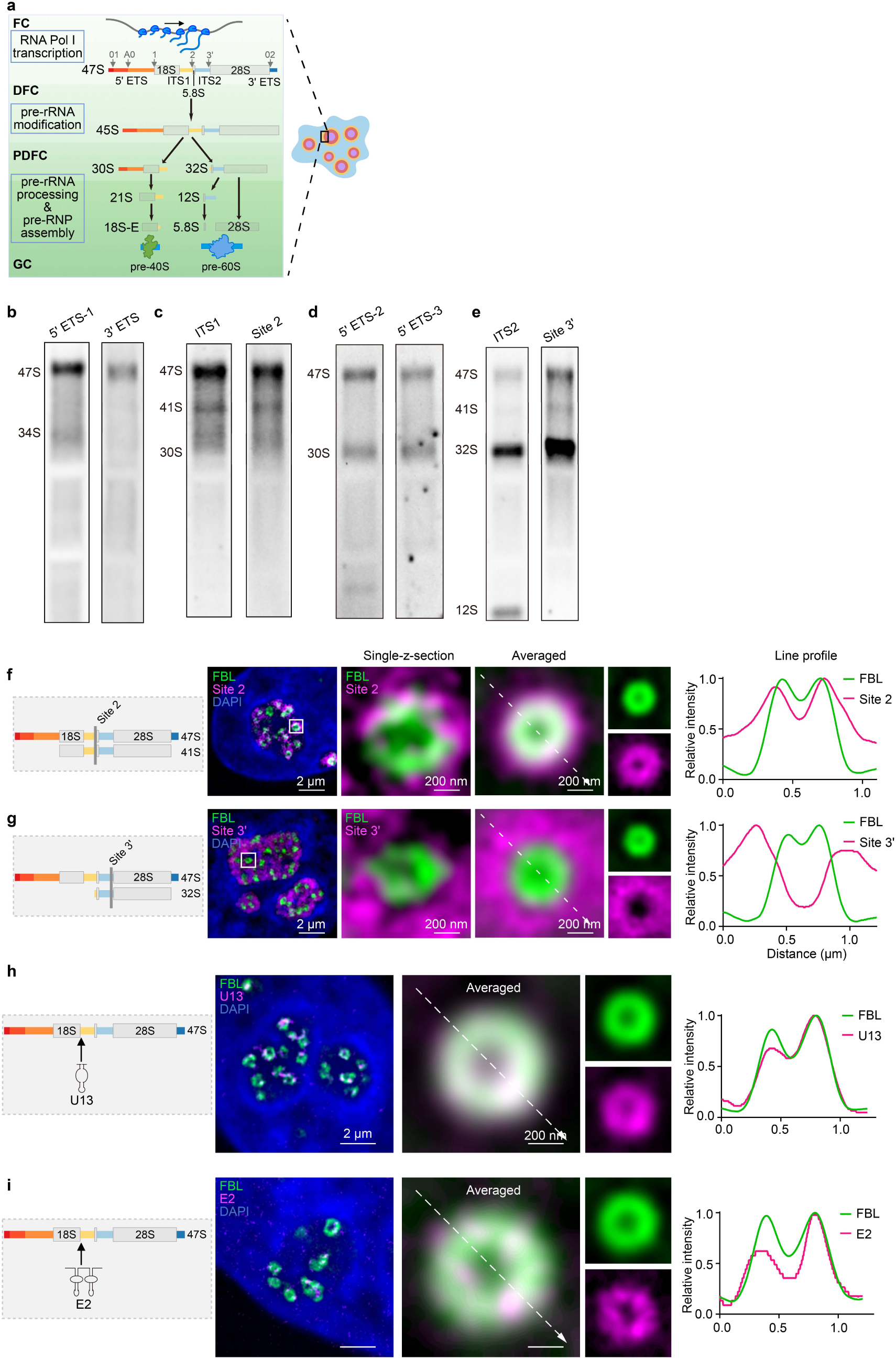
Sub-nucleolar localization of pre-rRNA intermediates by designed smFISH probes. **a**, Schematic of the highly organized subcompartments of nucleolus and the corresponding rRNA biogenesis events occurred in each sub-nucleolar region, including pre-rRNA transcription, modification, processing and the pre-rRNA ribonucleoprotein complex assembly. **b,** NB confirmed the designed probes, 5’ ETS-1 and 3’ ETS, mainly recognize the full length 47S pre-rRNA. See images in Fig. 1a. **c,** The ITS1-targeted probe sets, used for smFISH imaging, recognize the early state pre-rRNA, including mostly 47S pre-rRNA, and some 41S pre-rRNA. See images in Fig. 1c and panel **(f)**. **d,** The 5’ ETS-targeted probe sets, used for smFISH imaging, recognize the early state pre-rRNA and 18S rRNA precursors, including 47S and 30S pre-rRNA. See images in Fig. 1d. **e,** The ITS2-targeted probe sets, used for imaging, recognize 5.8S/28S rRNA precursors. The probe ITS2 used in Fig. 1e mainly recognizes the 32S and 12S pre-rRNA. The probe spanning site 3’ used in panel **(g)** mainly recognizes the 32S pre-rRNA. **f,** The Site 2 probe detected signal is retained within the DFC – PDFC region. Left, schematic of the localization of smFISH probe spanning the Site 2. Middle, representative single-sliced and averaged SIM images of the Site 2-detected 47S pre-rRNA (magenta) and FBL (green) in HeLa cells. Right, relative intensities of site 2 signals and FBL with radial position in the nucleolus. The number of DFC units (n) in the average images is 50 from 17 cells. **g,** The Site 3’-detected LSU pre-rRNA is mainly localized to the PDFC-GC region. Left, schematic of the localization of smFISH probe spanning the Site 3’. Middle, representative single-sliced and averaged SIM images of the Site 3’ detected 32S pre-rRNA (magenta) and FBL (green) in HeLa cells. Right, relative intensities of site 3’ signals and FBL with radial position in the nucleolus. The number of DFC units (n) in the average images is 50 from 17 cells. **h,** U13 snoRNA that binds the 3’ end of 18S pre-rRNA is mainly localized to the DFC. Left, schematic depicting the targeted position of U13 snoRNA on the pre-rRNA. Middle, representative single-sliced and averaged SIM images of U13 snoRNA (magenta) and FBL (green) in HeLa cells. Right, relative intensities of U13 snoRNA signals and FBL with radial position in the nucleolus. The number of DFC units (n) in the average images is 53 from 10 cells. **i,** E2 snoRNA that binds the 3’ end of 18S pre-rRNA is largely localized to the DFC. Left, schematic depicting the targeted position of E2 snoRNA on the pre-rRNA. Middle, representative single-sliced and averaged SIM images of E2 snoRNA (magenta) and FBL (green) in HeLa cells. Right, relative intensities of E2 snoRNA signals and FBL with radial position in the nucleolus. The number of DFC units (n) in the average images is 54 from 13 cells.

**Extended Data Figure 2.**
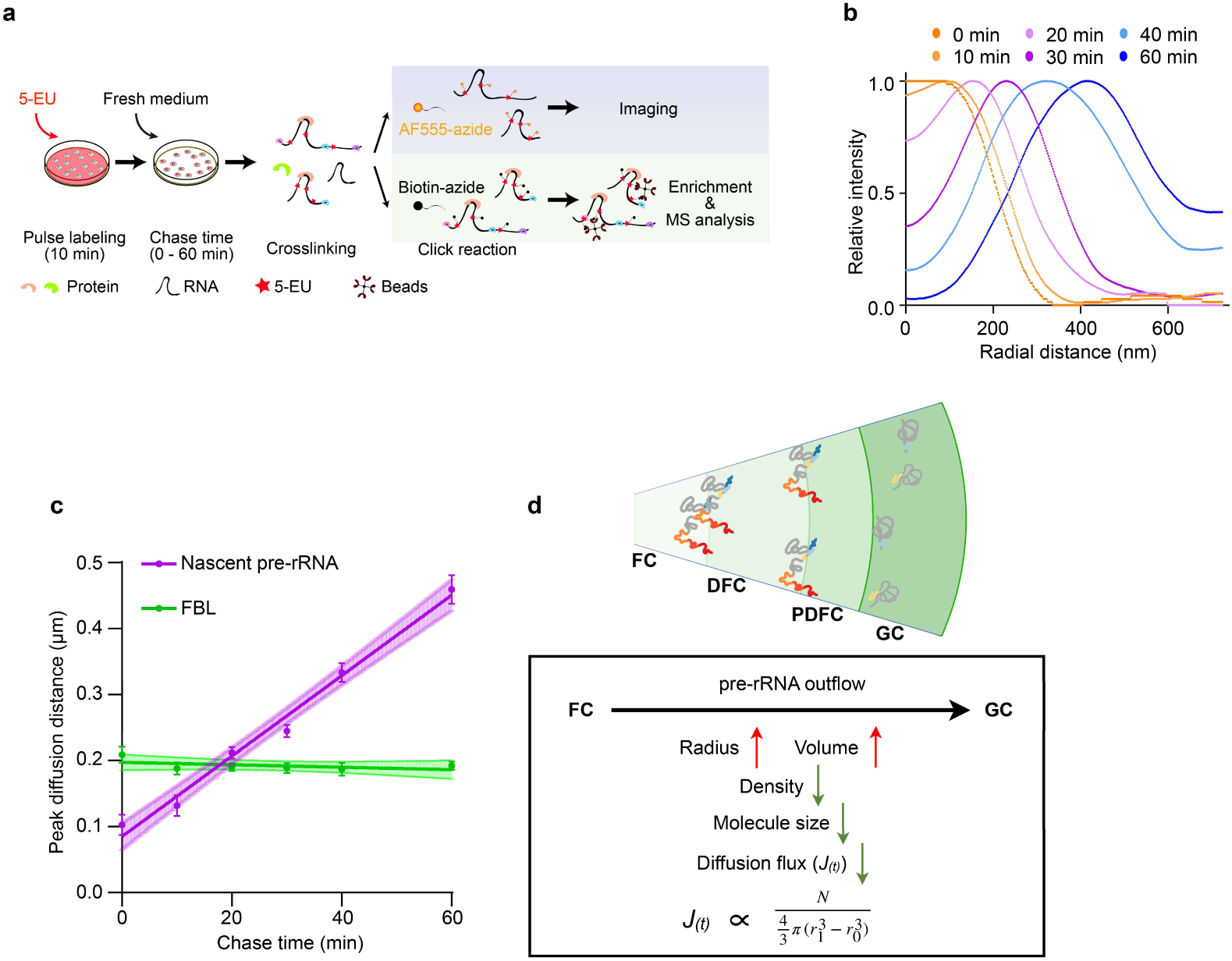
Characterization of the kinetics of nascent pre-rRNAs. **a**, The workflow of pulse-chase labeling experiment combined with super-resolution imaging and mass spectrometric (MS) analysis to examine the spatiotemporal distribution of SSU and LSU pre-rRNAs. Imaging measures the dynamics of pre-rRNAs flow in the nucleolus, while MS identifies nucleolar proteins associated with different stages of pre-rRNAs at indicated pulse-chase time points. **b,** Relative pre-rRNAs intensities with radial position from the FC center over time. Quantification of pre-rRNA intensities from averaged images in Fig. 1h, showing a gradient outflux distribution of nascent rRNAs across nucleoli over time. **c,** The peak diffusion distance of nascent pre-rRNAs over time (magenta), and the maximum intensity distance of FBL, which is constantly localized to the DFC, over time was used as a control (green). The line is fitted with linear regression with images of DFC units used for averaging in Fig. 1d. Data are mean ± s.e.m. n = 20 DFC units. **d,** Schematic of the pre-rRNA flow in nucleolar sub-compartments. Accompanying with the outward flow, pre-rRNAs are distributed in an increased 3D space, thus with a decreased spatial density. The relative diffusion flux is proportional to the density under the constant linear flow rate (calculated in panel **(c)**).

**Extended Data Figure 3.**
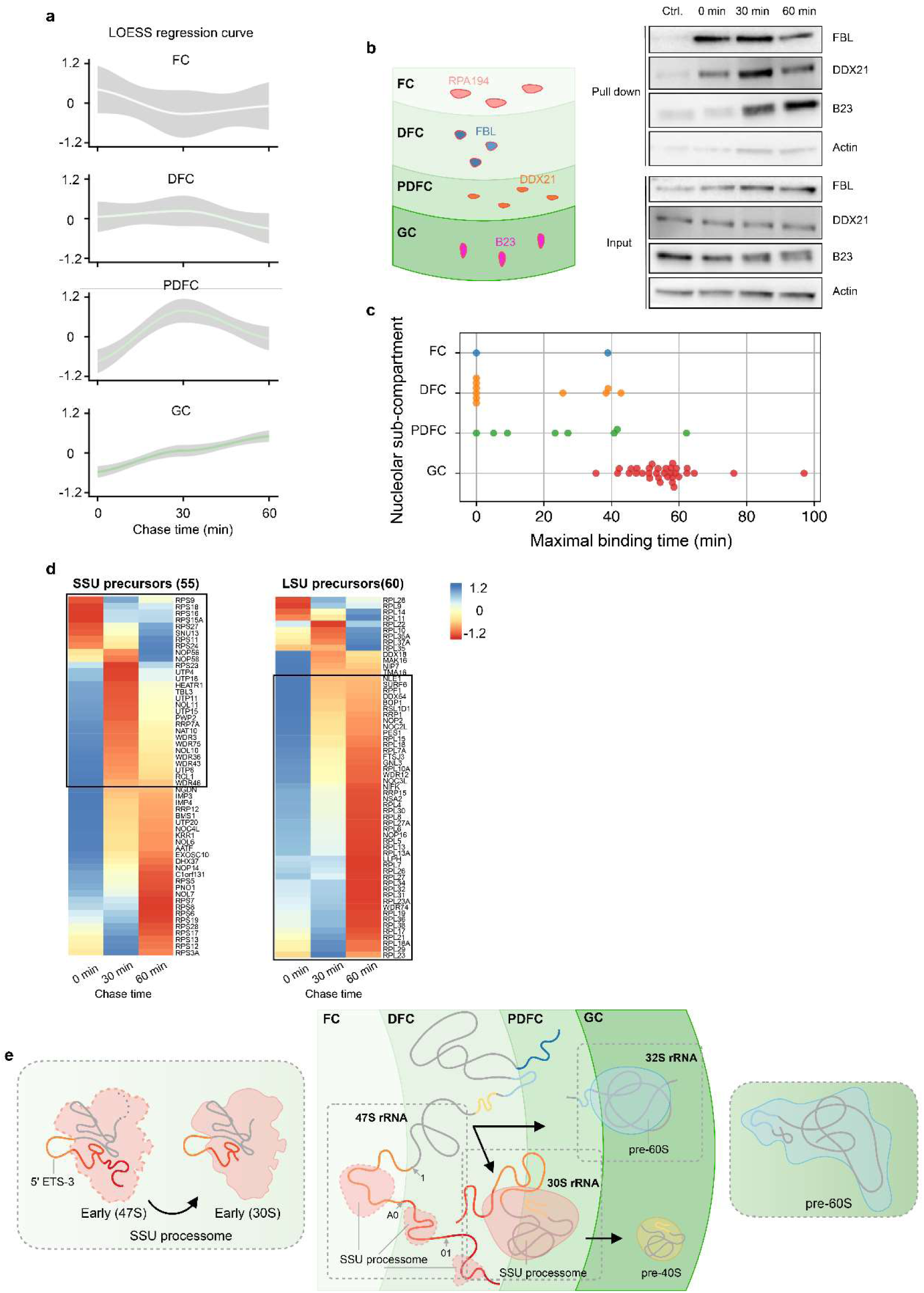
Characterization of the nascent pre-rRNA associated nucleolar proteins. **a**, The dynamic binding intensities of proteins in distinct nucleolar subdomains annotated in ^1^. The curve is fitted with LOESS regression. **b,** The dynamic enrichment of different nucleolar proteins in distinct nucleolar sub-domains over time upon pulse-chase labeling. Left, schematic of the nucleolar sub-domains with representative proteins, FC (RPA194), DFC (FBL), PDFC (DDX21), GC (B23) across different sub-domains. Right, Western blot (WB) analysis of proteins pull-downed by 5-EU-labeled nascent pre-rRNAs in **Extended Data** Fig. 2a. 5-EU nucleotide incorporation followed by biotin conjugation was used to precipitate proteins bound to nascent pre-rRNAs at different chase times. FBL is mostly enriched at the early chasing time points (10 min and 30 min), DDX21 is mostly enriched at the chasing 30 min, and B23 is mostly enriched at the 60 min chasing point. These dynamic distribution across the sub-nucleolar domains is correlated with the spatiotemporal distribution of nascent pre-rRNA shown in Fig. 1h. **c,** A recently published time-resolved mRNA interactome data using pulse-chase metabolic labeling with 4-thio-uridine (4sU)^28^ also revealed a similar outflow pattern of nucleolar protein localization. Consistent with the findings of this study, reported FC, DFC, and PDFC proteins bound to nascent rRNAs sequentially within the first 40-min chasing. And nascent rRNA-bound GC proteins presented after the 40-min time window. **d,** Quantities of SSU precursor proteins with peak enrichment time at 0min and 30min, LSU precursor proteins with peak enrichment time at 60min, both are marked by black box. The z-scores of log2 transformed protein LFQ intensities at each time point are in radial direction. **e,** The proposed model of the spatial distribution pattern of distinct pre-rRNA segments and the localization of the corresponding SSU and LSU pre-ribosomes.

**Extended Data Figure 4.**
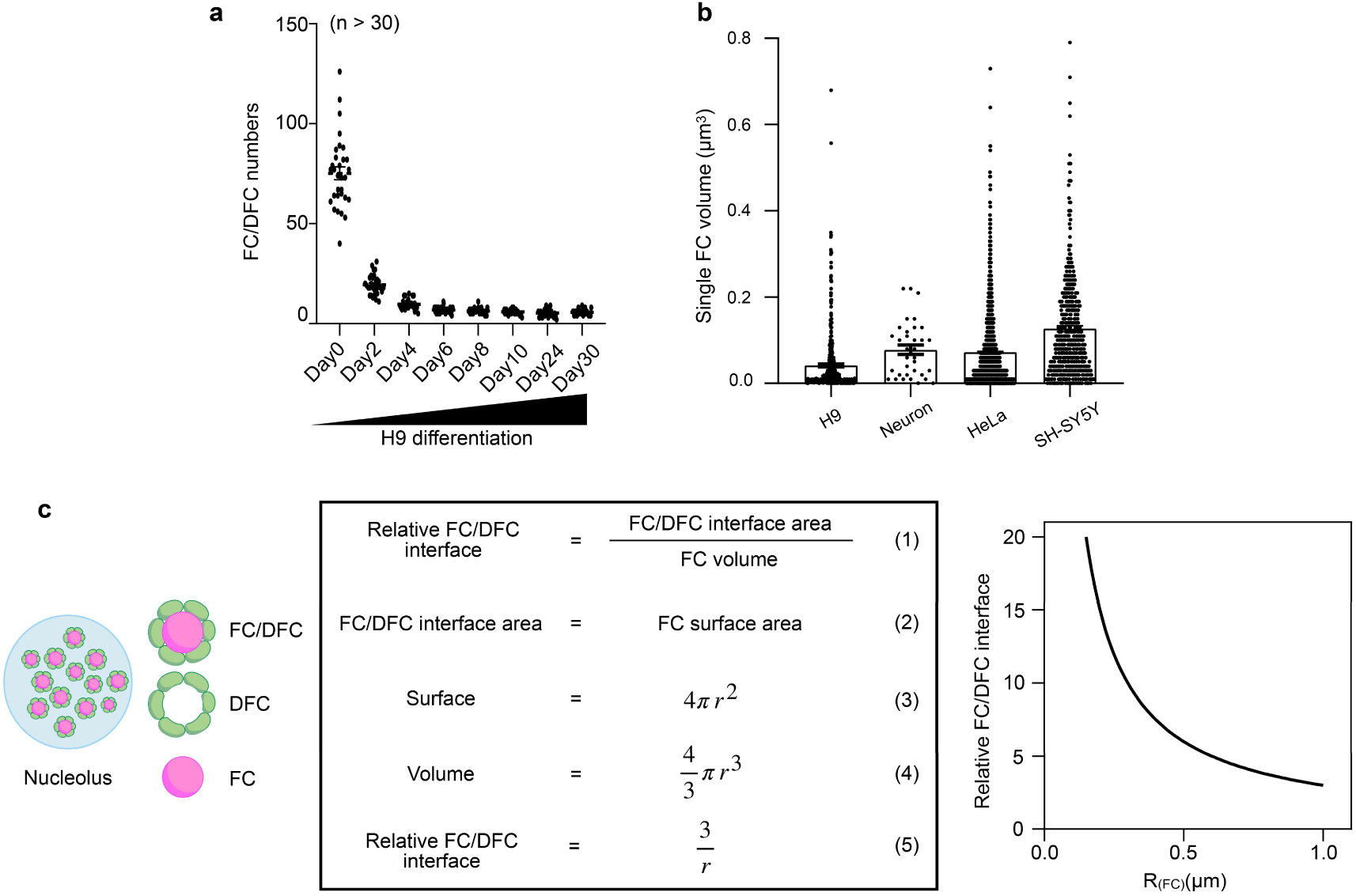
Analyses of the altered FC/DFC interface in morphologically different nucleoli from different cells. **a**, Statistics of FC/DFC units during H9 cells differentiation to neurons. Data are mean ± s.e.m. Number of cells (n) = 31, 31, 31, 47, 33, 31, 31, 32 at the sequential time points. **b,** The volume of the average FC in H9, H9-differentiated D30 neuron cells, HeLa and SH-SY5Y cells. Data are mean ± s.e.m. Data are from 4, 5, 30 and 30 cells, respectively. **c,** The theoretical relative FC/DFC interface in the spherical 3D is inversely proportional to the average FC radius. Left, the simplified model that assumes the FC and the DFC as nested spheres with the FC/DFC interface area presented as the FC surface. Middle, the calculation procedure that normalizes each FC/DFC interface area to individual FC volumes, obtaining the relative FC/DFC interface. Right, the plot of the theoretical function between the relative FC/DFC interface and the radius of the FC.

**Extended Data Figure 5.**
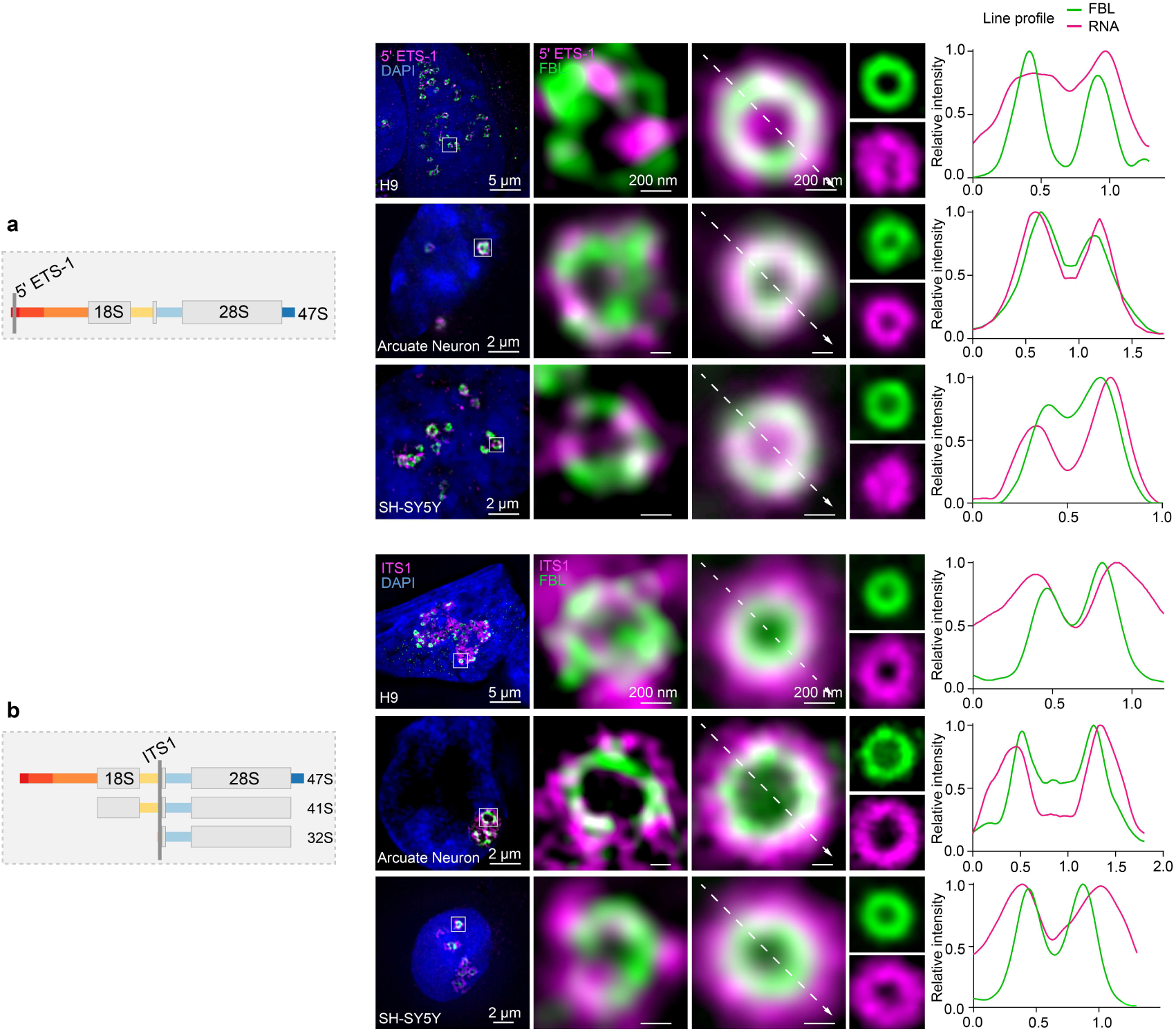
The spatial distribution of different segments of SSU pre-rRNAs in H9 cells, H9 differentiated neurons and SH-SY5Y cells. **a**, Localization of 5’ ETS-1 in the DFC region remain unchanged in different cell status. Left, schematic of the localization of the 5’ ETS-1 probe. Middle, representative single-sliced and averaged SIM images of 5’ ETS-1 (magenta) and FBL (green) in the indicated cell types. Right, the corresponding relative intensities of 5’ ETS-1 and FBL with radial position in the nucleolus. The number of DFC units (n) in the averaged images is more than 10. **b,** ITS1 localizes in the DFC-PDFC regions remain unchanged in different cell status. Left, schematic of the localization of the ITS1 probe. Middle, representative single-sliced and average SIM images of ITS1 (magenta) and FBL (green) in the indicated cell types. Right, the corresponding relative intensities of ITS1 and FBL with radial position in the nucleolus. The number of DFC units (n) in the averaged images is more than 10.

**Extended Data Figure 6.**
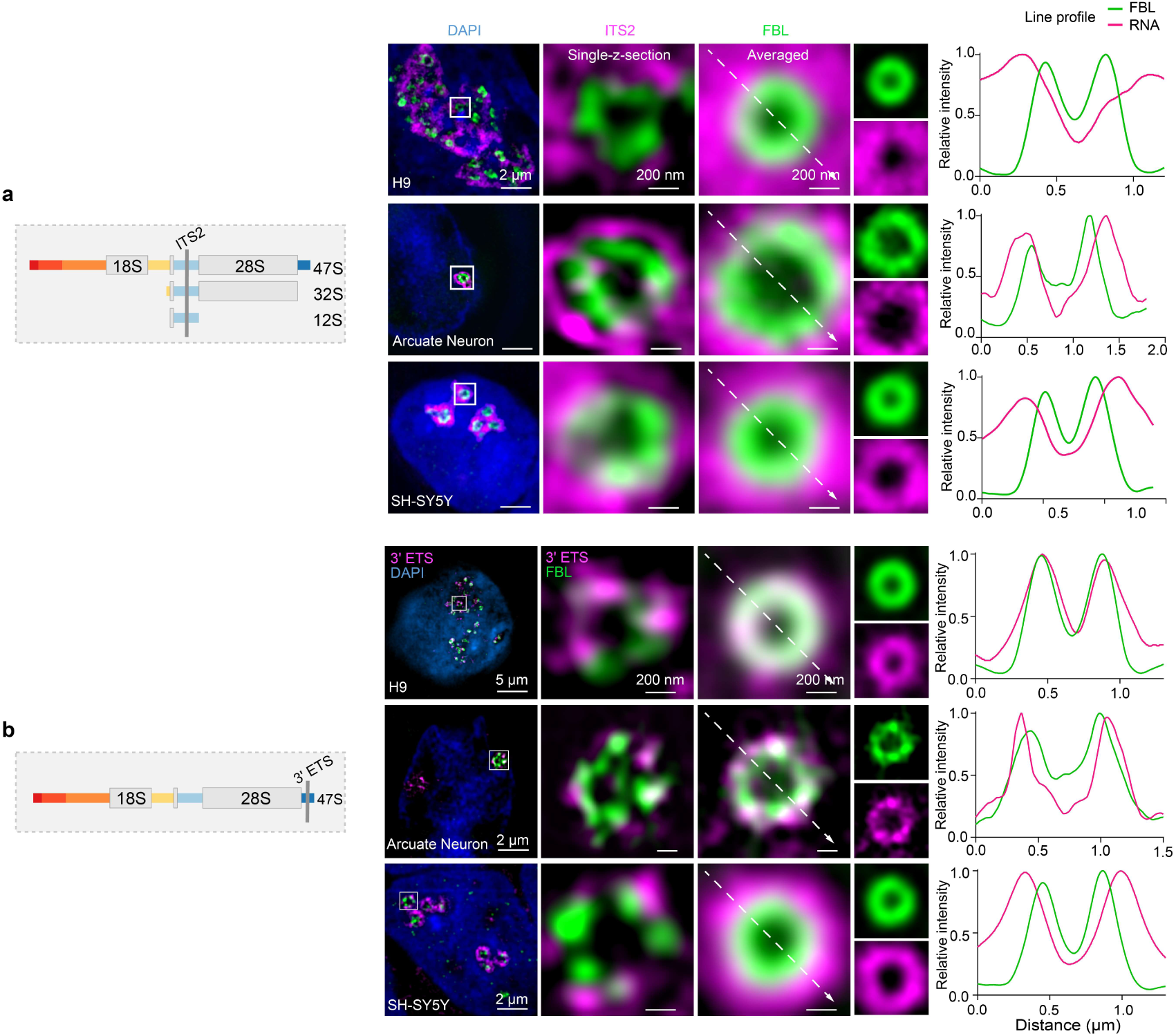
The spatial distribution of different segments of LSU pre-rRNAs in H9 cells, H9 differentiated neurons and SH-SY5Y cells. **a**, Localization of ITS2 in the PDFC-GC region remain unchanged in different cell status. Left, schematic of the localization of the ITS2 probe. Middle, representative single-sliced and average SIM images of ITS2 (magenta) and FBL (green). Right, the corresponding relative intensities of ITS2 and FBL with radial position in the nucleolus. The number of DFC units (n) in the averaged images is more than 10. **b,** Localization of 3’ ETS in the DFC-PDFC regions remain unchanged in different cell status. Left, schematic of the localization of the 3’ ETS probe. Middle, representative single-sliced and average SIM images of 3’ ETS (magenta) and FBL (green). Right, the corresponding relative intensities of 3’ ETS and FBL with radial position in the nucleolus. The number of DFC units (n) in the averaged images is more than 10.

**Extended Data Figure 7.**
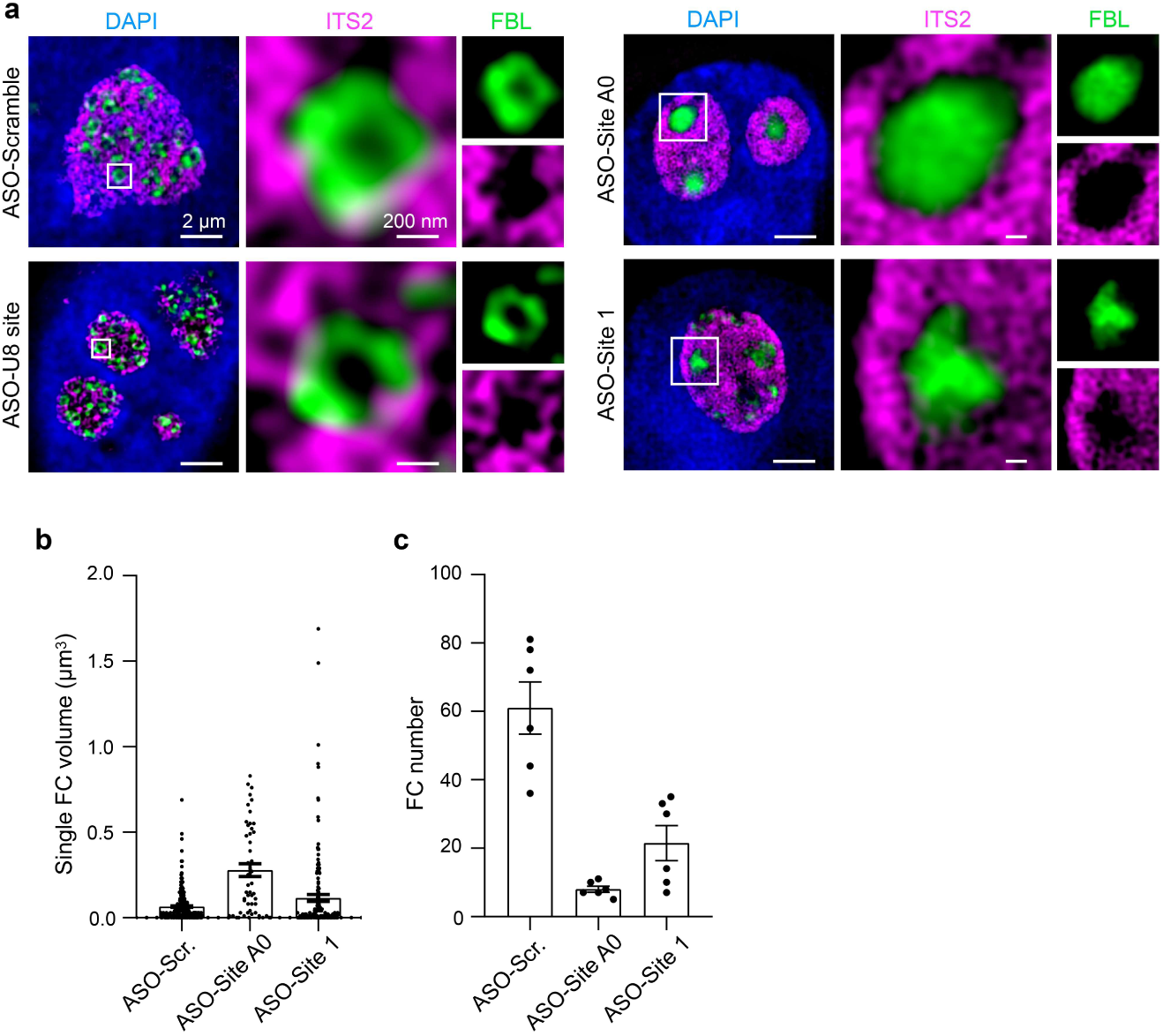
Re-organized FC/DFC sub-nucleolar structure upon disrupted SSU pre-rRNA processing. **a**, ITS2-detected LSU pre-rRNAs remain localized outside the DFC region marked by FBL after different ASOs treatment targeting 5’ ETS and the U8 snoRNA binding site. The stained ITS2 signal could be a marker of the GC region, like B23. **b,** The volume of the single FC in cells treated with ASO-Site A0 and ASO-Site1 that block SSU pre-rRNA processing was increased, compared to the ASO-Scr. Control treated HeLa cells. Data are mean ± s.e.m. (n) = 6 cells. **c,** The FC number in HeLa cells treated with ASO-Site A0 and ASO-Site1, respectively, was decreased, compared to that in cells treated with ASO-Scr. Data are mean ± s.e.m. Data are from 6 cells.

**Extended Data Figure 8.**
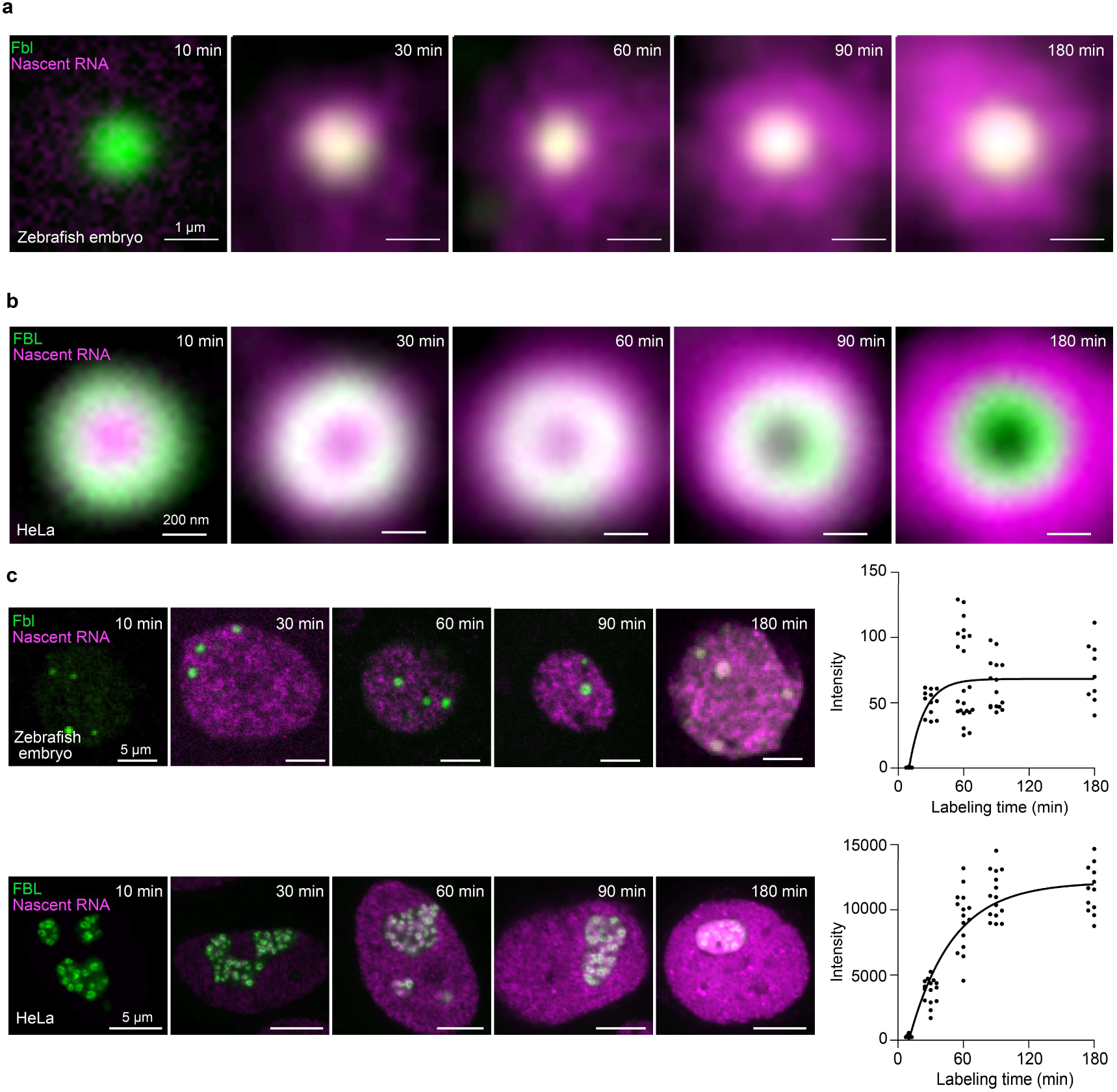
Distinct nascent RNA kinetics in zebrafish and HeLa nucleoli. **a**, Representative averaged images of nascent pre-rRNAs and Fbl at the sequential labeling time points in a cell obtained from zebrafish embryos. The signal of Fbl is a control to mark the F compartment in zebrafish nucleoli. See also Fig. 4d. **b,** Representative averaged images of nascent pre-rRNAs and FBL at sequential labeling time points in HeLa nucleoli. The signal of FBL is a control to mark the DFC. See also Fig. 4f. **c,** Representative whole-nucleus images of nascent pre-rRNAs and FBL, and statistics of nascent pre-rRNAs at the sequential labeling time points in zebrafish embryos and HeLa cells. Left, representative images of nascent rRNAs at different time points. Right, average nascent RNAs intensities in the nucleoplasm at sequential labeling time points, n>10. The curve is fitted using one-phase association. The intensity of Pol II-transcribed nascent RNAs (nucleoplasm distributed) did not decrease over the continuous labeling period, which excluded the possibility that 5-EU exhaustion caused the signal reduction of nascent pre-rRNA within the innermost FC region.

